# Cross-Propagative Graph Learning Reveals Spatial Tissue Domains in Multi-Modal Spatial Transcriptomics

**DOI:** 10.64898/2026.03.13.711582

**Authors:** Yin Guo, Songyan Liu, Zixuan Zhang, Shuqin Zhang, Limin Li

**Affiliations:** School of Mathematics and Statistics, Xi’an Jiaotong University, Xi’an, Shannxi, China; School of Mathematical Sciences, Fudan University, Shanghai, China

## Abstract

Spatial transcriptomics enables in situ characterization of tissue organization by jointly profiling gene expression profiles and spatial coordinates, with histological images as complementary contextual information. However, effectively integrating these heterogeneous modalities remains challenging due to differences in statistical properties and structural patterns. We propose st-Xprop, a cross-propagative graph network with dual-graph embedding coupling for spatial domain identification. st-Xprop constructs modality-specific graphs for gene expression and histological features, and performs alternating cross-modal propagation to explicitly model inter-modal heterogeneity while enabling complementary information exchange. Through dual-graph embedding coupling, the framework progressively learns a unified low-dimensional representation that integrates multimodal signals and preserves spatial coherence. Evaluations on multiple real spatial transcriptomics datasets demonstrate that st-Xprop consistently improves clustering accuracy and robustness, particularly in weak-signal or structurally complex regions, yielding spatial domains that are more stable and biologically meaningful.

## Introduction

Single-cell RNA sequencing has profoundly advanced the understanding of cellular transcriptional states, yet the loss of spatial context during tissue dissociation limits insight into tissue organization and spatially coordinated cell–cell interactions. Spatial transcriptomics (ST) addresses this limitation by capturing gene expression profiles within intact tissue architectures, enabling systematic investigation of spatial heterogeneity and functional organization in situ [43]. Recent technological developments have produced a diverse landscape of ST platforms, including imaging-based methods such as seqFISH [13], osmFISH [9], and MERFISH [58], as well as sequencing-based approaches such as 10x Visium [21], Slide-seq [36], and Stereo-seq [6]. These platforms differ substantially in spatial resolution, transcriptome coverage, and measurement noise, underscoring the need for robust computational methods to extract biologically meaningful tissue organization.

A core analytical task in spatial transcriptomics is the identification of spatial domains, i.e. contiguous tissue regions characterized by coherent transcriptional and histological features. Accurate spatial domain delineation provides a foundation for downstream analyses, including tissue annotation, developmental trajectory inference, and disease stratification. Early approaches largely adapted non-spatial clustering methods to ST data, relying exclusively on gene expression profiles and often producing spatially fragmented clusters that poorly reflect tissue architecture [44, 57]. Subsequent methods explicitly incorporated spatial coordinates, substantially improving spatial coherence by modeling neighborhood dependencies through probabilistic frameworks or graph-based neural networks, such as SpaGCN [20], STAGATE [10] and SEDR [53], thereby improving spatial coherence in domain identification. Despite these advances, most existing approaches remain focused on transcriptomic signals and geometric proximity.

Importantly, modern ST platforms such as 10x Visium routinely provide high-resolution histological images co-registered with gene expression measurements, offering an additional and complementary view of tissue structure. Histological images encode rich morphological information, including cues related to cellular density, tissue boundaries, and architectural motifs that are not directly observable from transcriptomic data alone. However, integrating histological and transcriptomic modalities poses a fundamental challenge due to their significant heterogeneity. Gene expression data are high-dimensional, sparse, and noisy, whereas histological images are dense, continuous, and characterized by complex spatial semantics. Simple feature concatenation or static fusion strategies fail to account for these differences and may obscure informative modality-specific structures rather than enhancing them. Several recent methods have attempted to leverage histological information for spatial domain identification by treating image-derived features as auxiliary inputs [20, 35]. While these approaches improve spatial smoothness in certain settings, histological information is typically incorporated in a static and secondary manner, without independent modeling of morphological structure or adaptive cross-modal interaction. As a result, these methods have limited capacity to capture complex and context-dependent relationships between tissue morphology and molecular states.

Here, we introduce st-Xprop, a cross-propagative graph network with dual-graph embedding coupling for spatial domain identification in spatial transcriptomics. st-Xprop explicitly models transcriptomic and histological modalities as two parallel yet interacting graph structures. A spatial graph encodes physical neighborhood relationships, while a histological graph captures morphological similarity derived from tissue images. The two graphs share a common set of nodes and take gene expression features as initial node attributes, while preserve modality-specific adjacency structures, enabling independent representation learning for each modality. To integrate multimodal information, st-Xprop employs a cross-propagative message-passing mechanism that exchanges information iteratively between the spatial and histological graphs. Learnable modality weights dynamically balance the contribution of each graph during propagation, allowing the model to adaptively calibrate spatial continuity and morphological boundary preservation. Through dual-graph embedding coupling and gene expression data fusion, st-Xprop progressively learns a unified low-dimensional representation that integrates complementary multimodal signals. We demonstrated the effectiveness of st-Xprop across multiple spatial transcriptomics datasets. st-Xprop consistently improves spatial domain identification accuracy and robustness compared to existing methods, particularly in regions with weak transcriptional signals or complex tissue architecture. The resulting spatial domains exhibit strong concordance with known anatomical structures and cell-type distributions, highlighting the significance of explicitly modeling cross-modal heterogeneity. Together, our results demonstrate that cross-propagative graph networks provide an effective and flexible framework for multi-modal integration in spatial transcriptomics, offering a robust foundation for downstream spatial analyses.

## Results

### Method Overview

st-Xprop is a cross-propagative graph network with dual-graph embedding coupling for spatial domain identification in spatial transcriptomics. The framework integrates gene expression, spatial coordinates, and histological image information by modeling transcriptomic and morphological signals as two complementary graph structures with shared nodes. A spatial graph encodes physical neighborhood relationships between spots, while a histological graph captures tissue morphology using image-derived features. Specially, st-Xprop introduces a cross-propagative message-passing mechanism that enables exchanging information flow between the spatial and histological graphs. Through learnable modality-specific weights, the model dynamically balances spatial smoothness and morphological boundary information during representation learning. By coupling embeddings across the two graphs and gene expression, st-Xprop learns a unified low-dimensional representation that preserves transcriptional patterns, spatial coherence, and histological structure. These coupled embeddings are subsequently used to identify coherent and biologically interpretable spatial domains and for other downstream analyses. The overall framework is summarized in Fig. 1.

**Figure 1.**
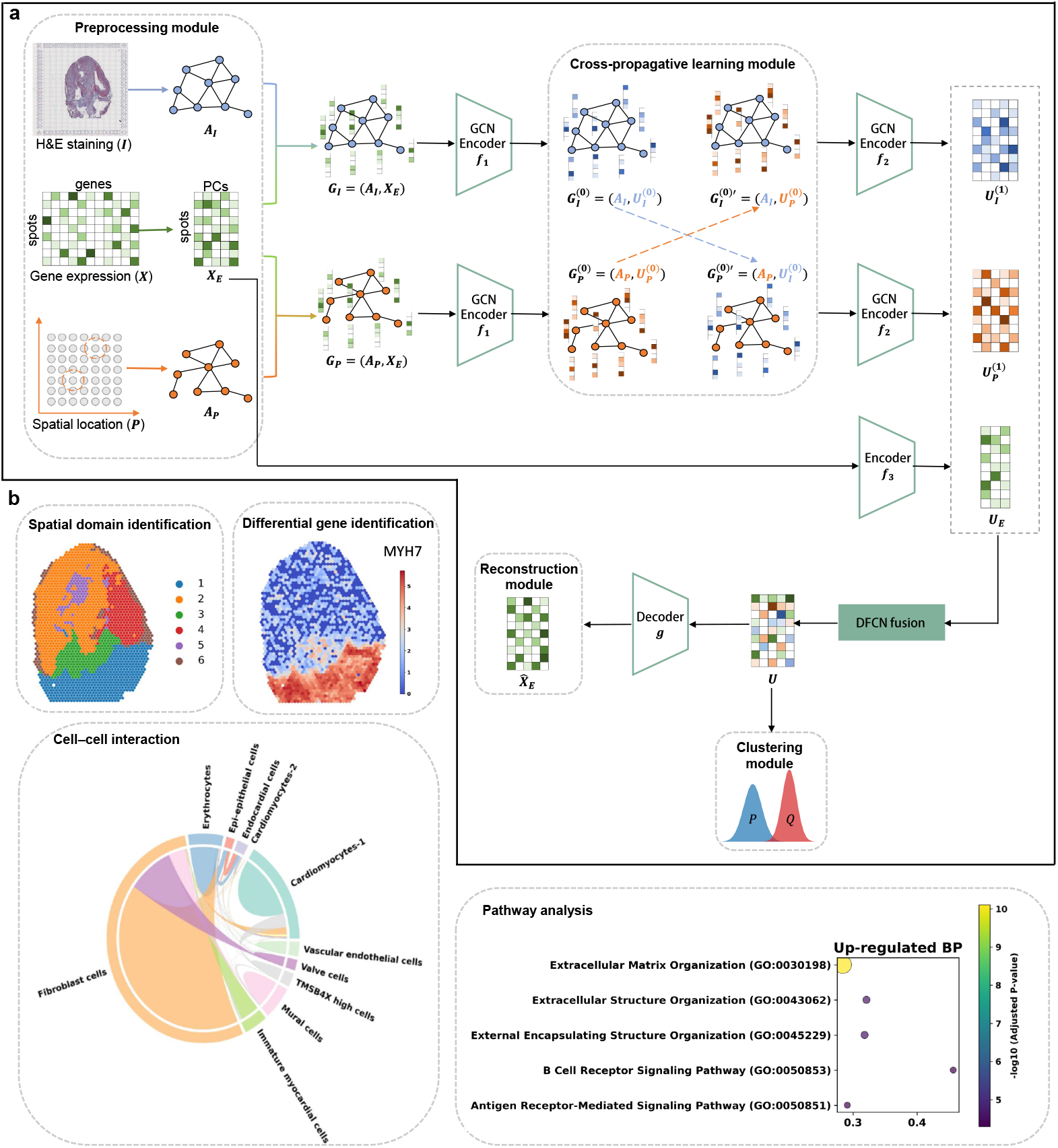
Overview of st-Xprop. (a) Framework overview. st-Xprop integrates gene expression **X**_*E*_, spatial coordinates **P**, and histological images **I** via dual-graph embedding coupling. A spatial graph **G**_*P*_ = (**A**_*P*_, **X**_*E*_) captures neighborhood structure, and a histological graph **G**_*I*_ = (**A**_*I*_, **X**_*E*_) encodes tissue morphology. Cross-propagative message passing alternates information flow, producing unified embeddings **U** for spatial domain identification. (b) Downstream analysis. The learned embeddings can be used for spatial domain identification and downstream tasks, including domain clustering, differential expression analysis, inference of cell–cell communication, and pathway enrichment.

To benchmark the performance of st-Xprop, we compared it with seven representative state-of-the-art methods for spatial domain identification, including DeepGFT [41], GraphST [54], SEDR [53], SpaGCN [20], spCLUE [50], STAGATE [10], and stLearn [35]. Among these methods, SpaGCN and stLearn additionally incorporate histological image information for spatial modeling, whereas the others primarily rely on gene expression and spatial structure (Experimental Section).

### st-Xprop Enables Accurate Spatial Domain Identification of Cortical Layers in Human DLPFC

We applied st-Xprop to the publicly available 10x Visium human dorsolateral prefrontal cortex (DLPFC) dataset [31], which contains manual annotations of cortical layers L1–L6 and white matter (WM) across 12 tissue sections. Based on morphological similarities, the sections were grouped into three classes, and one representative slice from each group (151507, 151671, and 151673) is highlighted in Fig. 2a, with the remaining slices shown in Fig. S1.

**Figure 2.**
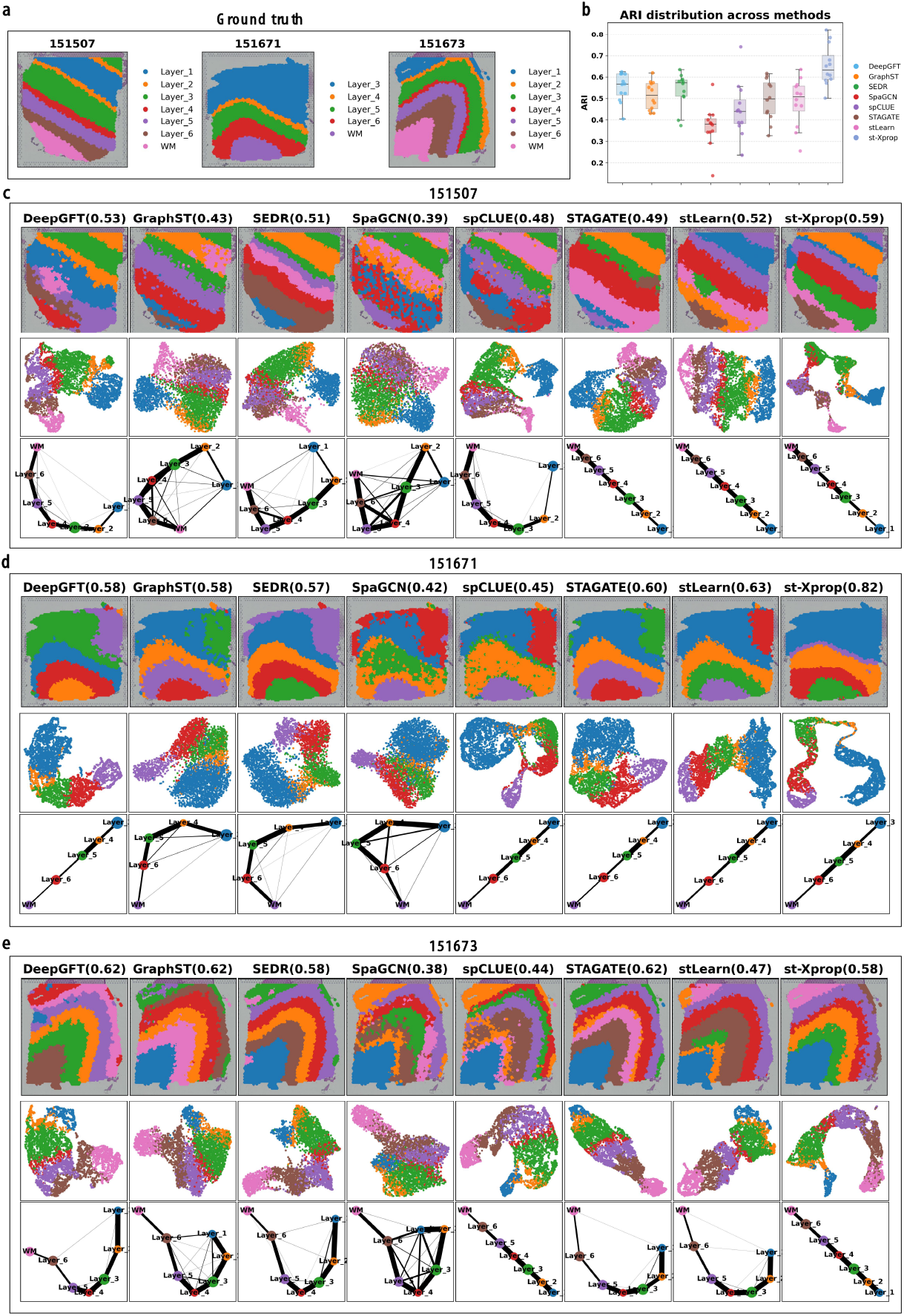
Spatial domain identification in human DLPFC. (a) Manual annotations of cortical layers in representative DLPFC slices 151507, 151671, and 151673. (b) Boxplots of ARI across all 12 slices. The center line, box edges, and whiskers indicate the median, first and third quartiles, and 1.5× interquartile range, respectively. (c–e) Comparison of spatial domain identification for slices 151507 (c), 151671 (d), and 151673 (e), with corresponding ARI scores, UMAP embeddings, and PAGA trajectories.

We first evaluated the overall performance of different methods across all 12 slices using the ARI metric (Fig. 2b). Among all slices, st-Xprop achieved the highest median ARI compared to other methods. Methods such as DeepGFT, GraphST, SEDR, STAGATE, and stLearn exhibited broadly similar performance, with most ARI values ranging between 0.5 and 0.6. In contrast, st-Xprop achieved ARI values above 0.6 on most slices, indicating competitive and generally improved clustering performance.

We next presented spatial clustering visualizations together with UMAP and PAGA representations based on the learned latent embeddings for the three representative slices (Fig. 2c–e). Across these slices, st-Xprop achieved the highest or comparable ARI scores relative to competing methods, while also providing more structurally coherent spatial domain assignments.

For slice 151507 (Fig. 2c), st-Xprop obtained the highest ARI and clearly resolved cortical layers L1–L3. In contrast, most competing methods failed to accurately identify layer L2, and merging it with adjacent layers L1 or L3. Although several methods captured the overall laminar organization, partial mixing between neighboring layers was observed in DeepGFT, SpaGCN, and spCLUE. In the UMAP embedding, st-Xprop yielded well-separated clusters corresponding to individual spatial domains, whereas other methods exhibited less distinct inter-cluster boundaries. The PAGA graph inferred by st-Xprop, as well as those from STAGATE and stLearn, revealed a largely linear backbone consistent with the known laminar ordering. However, other approaches introduced minor spurious branches or incorrect connections, suggesting less consistent global topology. These differences likely arise from the ability of st-Xprop to jointly integrate spatial adjacency and histological boundary information through cross-propagative learning, which helps preserve subtle layer transitions while maintaining spatial smoothness.

For slice 151671 (Fig. 2d), st-Xprop again achieved the highest ARI and correctly identified the cortical layers. By comparison, several competing methods misassigned layer L4 and further split layer L3 into multiple clusters, indicating over-segmentation and suboptimal alignment with anatomical boundaries. In both UMAP and PAGA representations, st-Xprop preserved a continuous and ordered structure consistent with the annotated laminar organization, whereas methods such as SEDR and SpaGCN exhibited less distinct cluster separation or trajectories with spurious branch connections.

In slice 151673 (Fig. 2e), the ARI of st-Xprop was slightly lower than that of DeepGFT, GraphST, and STAGATE. Although the grouping or separation of certain layers was not perfectly aligned with the discrete manual annotations, the spatial domains identified by st-Xprop exhibited smoother and more continuous patterns across the tissue. In contrast, the clustering results of the competing methods showed less spatial continuity, with occasional intra-layer fragmentation or less coherent cluster boundaries. As a result, their UMAP and PAGA representations were less consistent in preserving the linear laminar structure. In comparison, st-Xprop maintained clearer separation of major layers in UMAP and preserved a coherent linear backbone in PAGA that aligned with the known spatial ordering. These results indicate that st-Xprop better captures spatial continuity and structural coherence of cortical organization.

Overall, st-Xprop delineated spatial domains that are largely consistent with the annotated laminar structures in the DLPFC dataset. Similar spatial patterns were observed in the remaining nine slices (Fig. S1), further supporting the robustness of the method across slices.

### st-Xprop Identifies Anatomically Coherent Spatial Domains in the Mouse Brain

We next evaluated st-Xprop on 10x Visium mouse brain datasets, which exhibit finer and more heterogeneous spatial organization. Two representative slices were analyzed: a posterior sagittal slice (MBP) and a coronal slice (MBC), with anatomical regions annotated according to the Allen Mouse Brain Atlas [24] (Fig. 3a). For comparative analysis of clustering performance across methods, we focused on four representative anatomical regions across the two datasets, including three hippocampal formation (HPF) regions (square boxes 2, 3, and 4) and one cerebellar cortex (CBX) region (circular region 1). These regions were selected because they contain well-defined anatomical structures with varying spatial complexity, providing an informative setting for evaluating spatial domain identification. To further assess the ability of different methods to resolve fine-scale anatomical structures, we additionally examined substructures within region 4 of the hippocampal formation. Specifically, we focused on four well-characterized layers: CA1sp (field CA1, pyramidal layer), CA2sp (field CA2, pyramidal layer), CA3sp (field CA3, pyramidal layer), and DG-sg (dentate gyrus, granule cell layer), as indicated in the figure. In the absence of ground truth spatial domain labels, embeddings learned by different methods were clustered across multiple resolutions to assess the stability and granularity of spatial domain delineation. Specifically, Fig. 3b–c showed the spatial clustering results for MBP and MBC under a fixed setting of 18 clusters, while Fig. S2 and Fig. S3 present results across five resolutions (12, 16, 20, 22, and 25 clusters).

**Figure 3.**
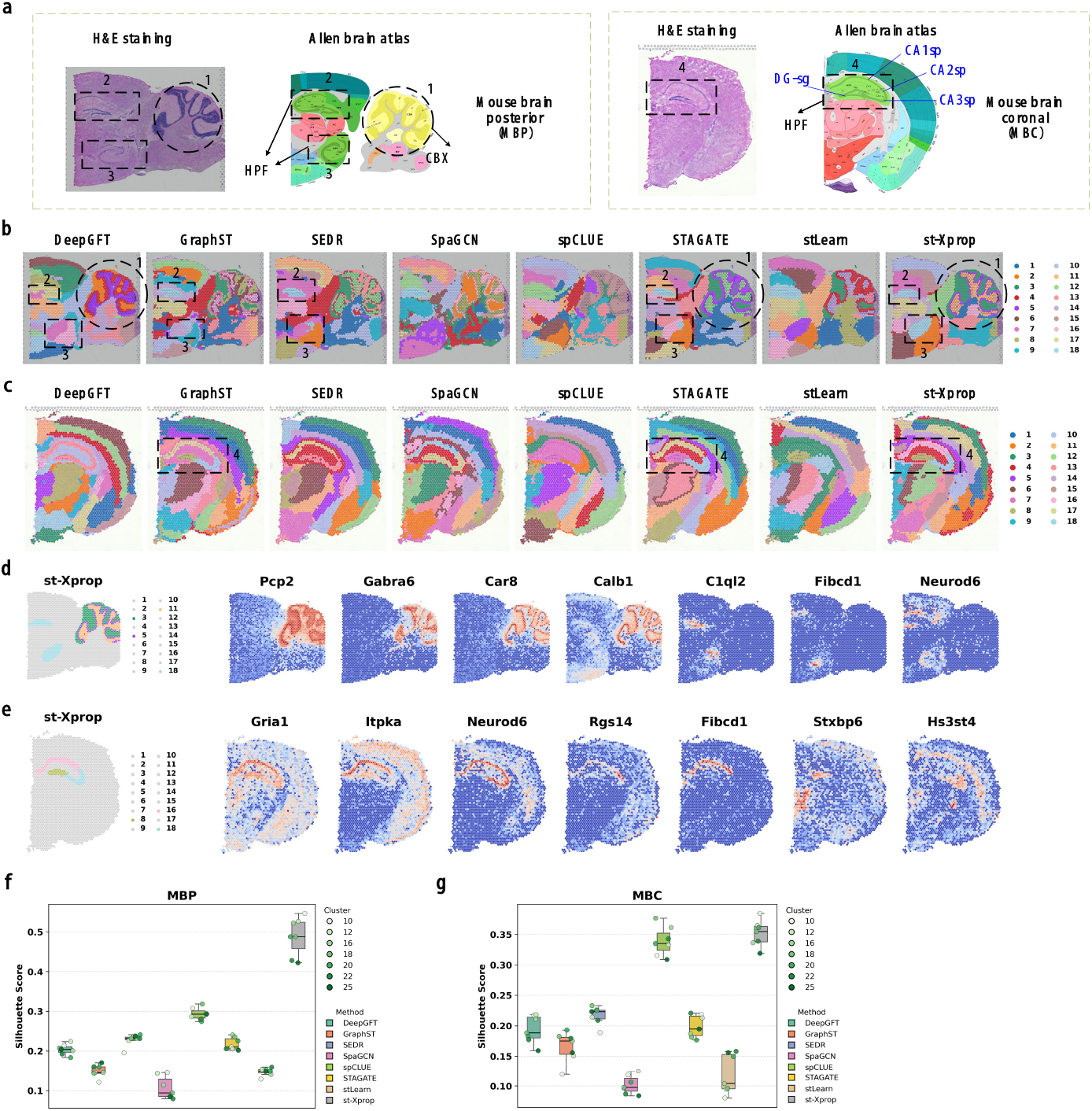
Spatial domain identification in mouse brain spatial transcriptomics. a) H&E-stained posterior sagittal (MBP) and coronal (MBC) mouse brain sections with anatomical references from the Allen Mouse Brain Atlas. Circles denote the cerebellar cortex (CBX), boxes highlight the hippocampal formation (HPF). b-c) Spatial domain clustering results obtained by st-Xprop and comparison methods on the MBP (b) and MBC (c) sections, shown at a clustering resolution of 18 domains. d-e) Spatial clustering results obtained by st-Xprop together with marker gene visualization for CBX and HPF regions in the MBP (d) and MBC (e) slices. f-g) Boxplots of silhouette scores across multiple clustering resolutions for the MBP (f) and MBC (g) sections. Center lines indicate medians; box limits represent the first and third quartiles; whiskers extend to 1.5× the interquartile range.

For the MBP slice (Fig. 3b and Fig. S2), most methods recovered the major anatomical regions across resolutions. The cluster partitions broadly corresponded to known anatomical compartments, although spatial continuity and boundary definition varied across methods. In the CBX region, DeepGFT, SEDR, STAGATE, and st-Xprop generated relatively continuous and well-delimited domains. Among them, st-Xprop exhibited the most spatially coherent partition, with smoother intra-region organization and clearer boundaries relative to adjacent regions. In addition, the identification of the hippocampal formation (HPF) showed substantial variation across methods. STAGATE and st-Xprop consistently grouped the two spatially separated HPF regions into a single coherent domain across clustering resolutions, indicating strong cross-space consistency. In contrast, DeepGFT, GraphST, SEDR, and SpaGCN identified HPF only at specific resolutions, while SpaGCN and stLearn frequently split the two HPF regions into separate clusters. GraphST and SEDR occasionally merged portions of HPF with neighboring regions, resulting in incomplete recovery of the anatomical structure. These differences reflect varying levels of stability in cross-region aggregation.

Across the entire tissue section, st-Xprop produced partitions with stronger spatial continuity and clearer boundaries compared to other methods. This qualitative observation was supported quantitatively by silhouette scores (Fig. 3f), where st-Xprop consistently achieved higher values across all clustering resolutions, indicating improved intra-cluster compactness and inter-cluster separation.

To further assess biological relevance, marker genes were examined (Fig. 3d). Cerebellar markers Pcp2, Gabra6, Car8, and Calb1 showed enriched expression within the CBX domains identified by st-Xprop (domain union 3, 5, and 11), while hippocampal-associated markers C1ql2, Fibcd1, and Neurod6 were localized within the HPF region (domain 18). The spatial distribution of these markers aligned closely with the st-Xprop partitions, supporting the anatomical fidelity of the inferred domains.

For the MBC slice (Fig. 3c and Fig. S3), most methods delineated the overall hippocampal formation across clustering resolutions, while their ability to resolve internal layers differed substantially. In the hippocampal substructure highlighted in region 4, st-Xprop clearly separated the DG-sg and CA1sp layers into distinct spatial domains, while grouping the anatomically adjacent CA2sp and CA3sp layers into a single coherent domain. This partitioning preserved the layered hippocampal organization and maintained spatial continuity with well-defined boundaries. DeepGFT, SEDR, and SpaGCN generally captured the overall HPF but merged the subregions into one or two coarse clusters, limiting representation of internal structural heterogeneity. GraphST and STAGATE revealed partial HPF subregions as cluster numbers increased, yet the recovered structures varied across resolutions, indicating reduced robustness. spCLUE and stLearn identified some substructures at higher resolutions but frequently mixed them with adjacent cortical areas, leading to less anatomically consistent partitions. These qualitative findings were supported by silhouette score analysis (Fig. 3g), where st-Xprop consistently achieved higher and more stable silhouette scores across clustering resolutions, indicating improved cluster compactness and separation.

Marker gene visualization further supported the anatomical consistency of the inferred domains (Fig. 3e). DG- and CA-associated markers, including Gria1, Itpka, Neurod6, Rgs14, Fibcd1, Stxbp6, and Hs3st4, exhibited subregion-specific enrichment patterns that were spatially aligned with the domains identified by st-Xprop (domain union 8, 16, and 18). The observed correspondence between marker localization and cluster assignment indicates that the inferred partitions capture biologically meaningful hippocampal substructures.

Collectively, these results indicate that st-Xprop yields fine-grained and stable spatial domain representations that are anatomically consistent in complex mouse brain tissues, with competitive performance in highly heterogeneous spatial contexts.

### st-Xprop Reveals Spatiotemporal Organization and Cell–cell Communication During Chicken Heart Development

We next evaluated st-Xprop on a 10x Visium chicken heart spatial transcriptomics dataset spanning four consecutive developmental stages (D4, D7, D10, and D14), for which manual spatial annotations are available [30]. As summarized in Fig. 4a–d, st-Xprop consistently achieved the highest ARI across all stages, whereas other methods showed lower values and greater variability. Spatial visualization further illustrated these differences. Across D4–D14, st-Xprop reconstructed anatomically defined heart regions with clear boundaries and spatial continuity, closely matching known developmental structures. In contrast, other methods frequently exhibited region mixing, partial loss of thin structures, or blurred boundaries between adjacent compartments.

**Figure 4.**
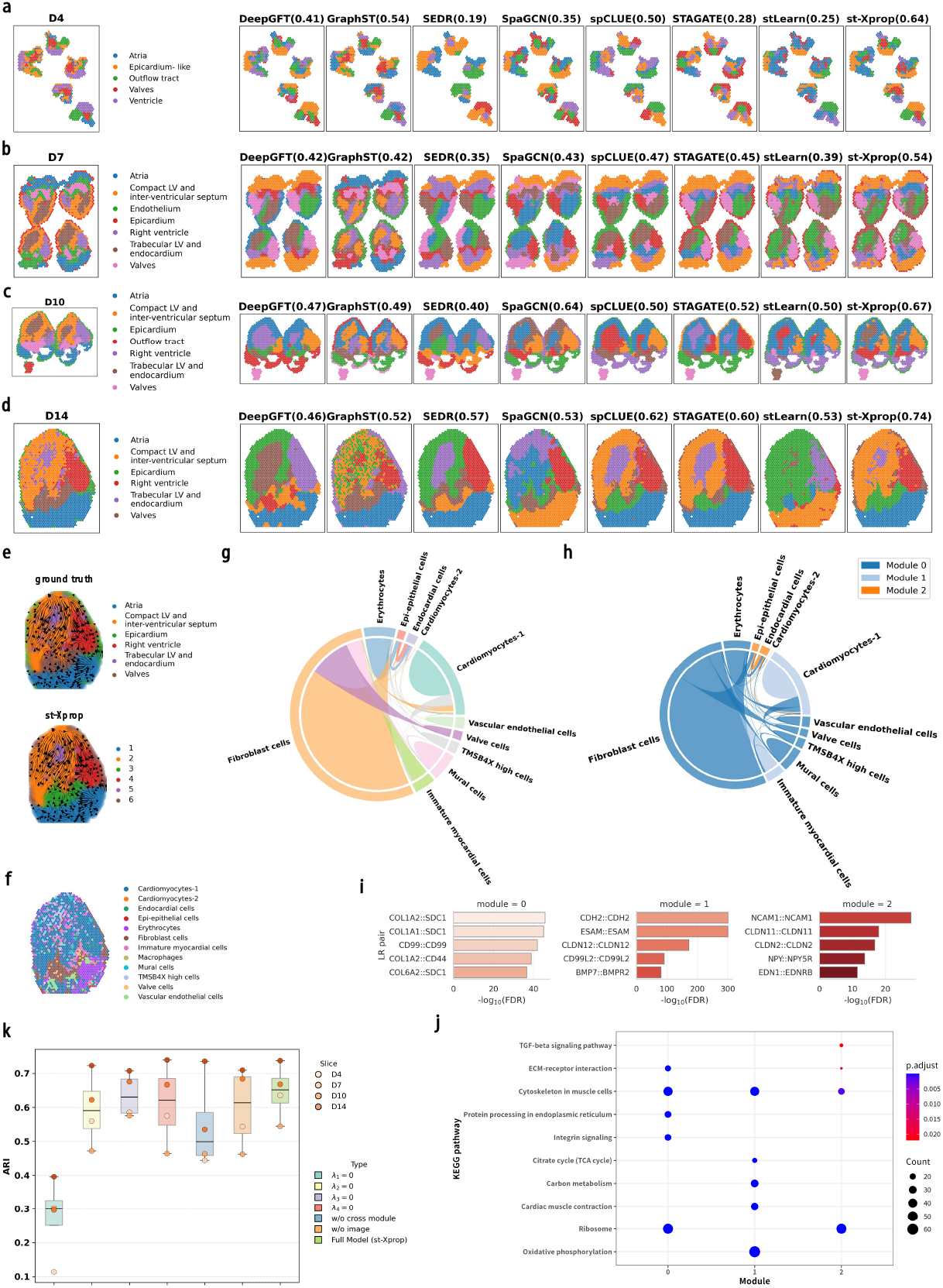
Spatial domain organization and cell–cell communication in the developing chicken heart. a–d) Spot-level manual annotations, spatial domain identification by different methods, and corresponding ARI scores across four developmental stages (D4, D7, D10, and D14). e) Spatial RNA velocity field for the D14 slice, colored by anatomical annotations or st-Xprop-identified spatial domains. f) Spatial distribution of major annotated cell types in the D14 slice. g) Cell–cell interaction (CCI) network inferred from ligand–receptor co-expression and spatial proximity based on st-Xprop spatial representations, and visualized as a chord diagram. h) Communication modules identified from the CCI network. i) Enriched ligand–receptor pairs within each communication module. j) KEGG pathway enrichment analysis of module-associated genes. k) Ablation analysis quantifying the contribution of individual st-Xprop modules to spatial domain identification, evaluated by ARI across developmental stages.

At D4 (Fig. 4a), st-Xprop separated the atrial and ventricular compartments and further delineated the outflow tract and valvular regions as distinct spatial domains. The inferred partitions were spatially continuous and aligned with the annotated anatomical layout, with clear boundaries between adjacent regions. In comparison, DeepGFT, GraphST, and spCLUE recovered the overall heart contour but showed incomplete separation of specific compartments: DeepGFT did not fully distinguish the outflow tract from the valves, GraphST partially mixed atrial and ventricular regions, and spCLUE failed to clearly resolve atrial and valvular domains. SpaGCN exhibited mixing across adjacent regions, while SEDR, STAGATE, and stLearn produced less distinct ventricular substructures, resulting in blurred internal boundaries.

At D7 (Fig. 4b), when the architecture becomes more complex, st-Xprop resolved multiple anatomically and functionally distinct subregions. Thin and spatially restricted structures such as valves and epicardium were delineated as spatially continuous domains with coherent boundaries. In comparison, DeepGFT, SEDR, SpaGCN, STAGATE, and stLearn frequently merged compact and trabecular left ventricular regions into a single domain. GraphST grouped the trabecular left ventricle together with the right ventricle, reducing ventricular subtype resolution. Additionally, spCLUE and SEDR did not clearly delineate thin-layer structures such as valves and epicardium, instead merging them with adjacent regions.

In the later stages D10 (Fig. 4c) and D14 (Fig. 4d), st-Xprop delineated trabecular and compact left ventricular compartments as distinct domains and preserved spatially continuous valve and epicardium regions with well-defined boundaries. By comparison, DeepGFT and SEDR did not reliably recover thin-layer structures such as the epicardium. Moreover, all comparative methods exhibited varying degrees of mixing between trabecular and compact left ventricular regions in both stages, leading to less distinct delineation of ventricular subtype organization. Across developmental stages, st-Xprop consistently recovered the expected spatial arrangement and morphology of major cardiac domains.

To assess whether the inferred spatial domains reflect coherent transcriptional dynamics, we computed RNA velocity fields on the D14 dataset using scVelo [3] and projected them onto tissue sections under both anatomical annotations and st-Xprop-defined clusterings (Fig. 4e). RNA velocity vectors exhibited spatially organized patterns, with consistent directional flow within st-Xprop domains and gradual changes across domain boundaries. Similar patterns were observed under anatomical annotations, indicating that the inferred domains are compatible with the underlying transcriptional dynamics.

In addition to anatomical annotations, spot-level major cell-type labels were available for this dataset. We visualized the spatial distribution of major cell types in the D14 slice (Fig. 4f) to examine whether the st-Xprop-inferred domains correspond to biologically meaningful cellular organization. Cardiomyocytes, TMSB4X-high cells, and a subset of immature myocardial cells were predominantly enriched in ventricular compartments, including the compact left ventricular and interventricular septum, right ventricle, and trabecular left ventricular with endocardium, corresponding to st-Xprop-defined domains 2, 4, and 5. These regions form a spatially continuous ventricular territory in the tissue section. Fibroblasts, valve-associated cells, and some immature myocardial cells were concentrated in the valve region (domain 3), whereas vascular endothelial cells and erythrocytes were primarily localized to the atrial region (domain 1). Epi-epithelial and endocardial cells, although less abundant, were also mainly observed near the atrial region. In addition, fibroblasts and erythrocytes showed enrichment in the epicardium (domain 6). Importantly, the spatial enrichment patterns of these major cell populations align closely with the st-Xprop-defined domains. This concordance indicates that the inferred spatial partitions reflect biologically coherent tissue compartments rather than arbitrary clusters, supporting the ability of st-Xprop to learn spatial representations that capture underlying cellular organization.

To further examine how these spatial patterns relate to intercellular communication, we constructed a global cell–cell interaction (CCI) network by integrating cell type labels, st-Xprop embeddings, and ligand–receptor co-expression information [8] (Experimental Section). The chord diagram (Fig. 4g) revealed a structured communication network inferred from ligand–receptor interactions. Fibroblasts occupied a central position in the network, receiving substantial incoming signals from erythrocytes and cardiomyocytes and exhibiting enriched outgoing interactions toward valve-associated cells, mural cells, and immature myocardial cells. Notably, fibroblasts were primarily localized in domain 3, a valve-associated region that lies adjacent to multiple neighboring domains. This spatial positioning likely contributes to their high network centrality, as interacting cell types are located in nearby tissue compartments. Cardiomyocytes, particularly the Cardiomyocytes-1 subset, displayed dominant outgoing signaling activity, including predicted interactions with TMSB4X-high cells. These interactions are consistent with their shared enrichment in ventricular domains (domains 2, 4, and 5), where these cell populations form spatially contiguous regions. Similarly, erythrocytes received prominent signals from epi-epithelial and endocardial cells, which are co-localized in the atrial region (domain 1), supporting the spatial plausibility of these inferred interactions. Together, these observations indicate that the inferred CCI patterns are strongly shaped by the spatial organization captured by st-Xprop, with interacting cell populations largely co-localized or positioned in adjacent tissue domains. This spatially coherent communication landscape further supports the ability of st-Xprop to recover biologically meaningful tissue organization. These inferred communication patterns are concordant with previous single-cell and developmental studies in human and murine hearts. For example, recent spatial and single-cell analyses have identified cardiac fibroblasts as high-centrality nodes within ligand–receptor networks, exhibiting extensive signaling interactions with cardiomyocytes [15, 28]. Developmental studies further demonstrate that resident cardiac fibroblasts arise from mesenchymal lineages through epithelial–mesenchymal transition (EMT) and endothelial–mesenchymal transition (EndMT) during embryogenesis [37], and that epicardial-derived mesenchymal cells subsequently differentiate into mural cells and fibroblasts during cardiac growth [40]. In addition, thymosin *β*4 (TMSB4X) has been shown to promote cardiomyocyte migration and survival [4], supporting the biological plausibility of the predicted interactions involving TMSB4X-high cells. Given the evolutionary conservation of key cardiac developmental pathways across vertebrates, these findings in chicken heart tissue align with established mechanisms reported in human and mouse hearts.

To further characterize inter-domain communication, we applied Leiden clustering to the CCI network and identified three major modules (Fig. 4h), each exhibiting clear cellular and spatial coherence: (i) a stroma–vascular module (module 0) enriched for fibroblasts, vascular endothelial, mural, and valve-associated cells; (ii) a cardiomyocyte-centered module (module 1) dominated by muscle and immune-related populations; and (iii) a luminal module (module 2) composed primarily of epithelial and endocardial cells. We next examined ligand–receptor (LR) interactions and associated gene programs within each module (Experimental Section; Fig. 4i–j). The stroma–vascular module (module 0) showed strong enrichment for extracellular matrix (ECM)-related interactions, including collagen–proteoglycan pairs (e.g., COL1A1/2–SDC1) and collagen–CD44, with KEGG pathway analysis highlighting ECM–receptor interaction and integrin signaling pathways. The cardiomyocyte module (module 1) was dominated by homophilic adhesion and developmental signaling pairs, such as CDH2–CDH2, CD99L2–CD99L2, and BMP7–BMPR2, with corresponding enrichment in pathways related to cytoskeletal organization and cardiac contraction. The luminal module (module 2) exhibited enrichment for junctional adhesion and paracrine signaling interactions, including NCAM1–NCAM1, claudin family pairs, and EDN1–EDNRB signaling. These interactions were associated with pathways involved in ECM–receptor interaction, cytoskeletal organization, and TGF-beta signaling. Together, these analyses indicate that the inferred spatial domains support structured and functionally coherent communication modules during heart development.

We next performed ablation studies to assess the contributions of individual components of st-Xprop across all developmental stages (Fig. 4k). Ablations included: (i) setting each individual loss weights to zero; (ii) removing the cross-propagative learning module; and (iii) omitting histological image data, using only gene expression and spatial coordinates. Results indicated that exclusion of the reconstruction loss ℒ_rec_ (*λ*_1_ = 0) resulted in a marked reduction in ARI, underscoring the importance of learning robust low-dimensional representations. Removal of the cross-propagative learning module similarly impaired performance, highlighting its role in integrating gene expression with histological features. Omitting histological image information and relying only on gene expression and spatial coordinates also led to decreased accuracy, indicating that morphological cues provide complementary information for spatial domain identification. In all cases, the full st-Xprop model achieved the highest and most stable ARI across stages.

Collectively, these results show that st-Xprop supports accurate spatial domain identification across developmental stages and enables coherent analysis of transcriptional dynamics and intercellular communication within a unified spatial framework.

### st-Xprop Identifies Spatially Organized Tumor Regions in Breast Cancer

We next assessed the performance and generalizability of st-Xprop in a cancer setting using a publicly available 10x Visium breast cancer (BRCA) dataset. This tissue section was manually annotated by pathologists into 20 subregions, grouped into four histologically defined classes (Fig. 5a–b). Spatial domains inferred by different methods were compared against these annotations (Fig. 5c). st-Xprop achieved the highest agreement with manual labels (ARI = 0.65), and its inferred domains exhibited smoother boundaries that more closely followed histological structures. In contrast, several methods failed to maintain the integrity of specific annotated regions. Most comparative methods, except stLearn, generated fragmented and spatially discontinuous domains across the tissue section, which likely contributed to their lower ARI values. For example, in the upper-right Tumor edge 6 region and the lower-right IDC 2 region, DeepGFT, GraphST, SEDR, spCLUE, and STAGATE partitioned these regions into multiple discontinuous domains rather than preserving them as coherent spatial units. Notably, although st-Xprop-defined domain 3 appeared spatially discontinuous, these regions corresponded entirely to the annotated Tumor edge, indicating that the model captured histologically coherent features that are not strictly defined by spatial contiguity.

**Figure 5.**
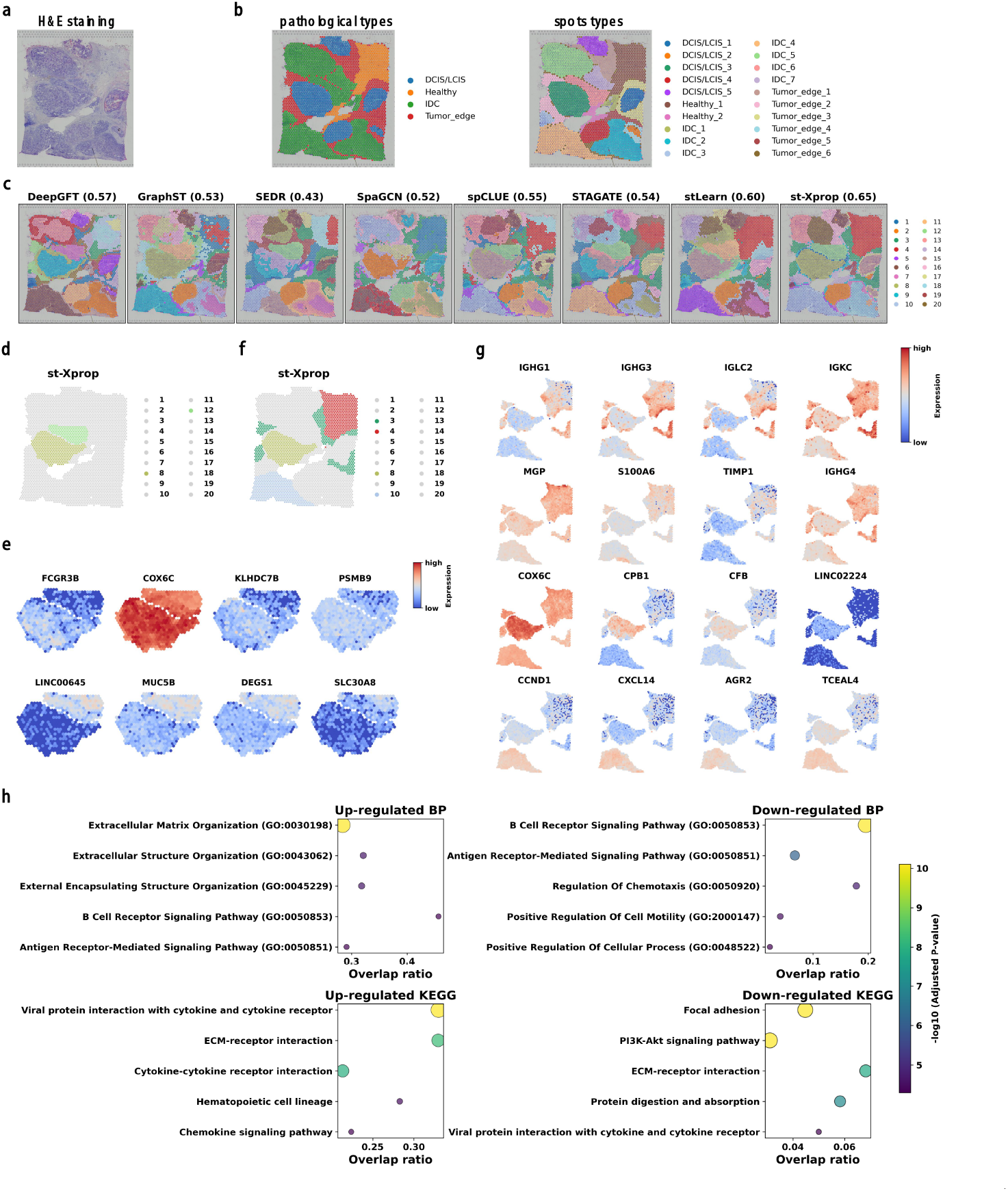
Spatial domain identification results on human breast cancer tissue. a) Histological image of the breast cancer tissue section. b) Manual spot-level annotations with pathological subtypes. c) Spatial domain identification by st-Xprop and baseline methods, with corresponding adjusted Rand index (ARI) scores evaluated against manual annotations. d) Spatial distribution of st-Xprop-identified domains 8 and 12 within the DCIS/LCIS–IDC composite region. e) Differentially expressed genes between domains 8 and 12 (top: domain 8; bottom: domain 12). f) Spatial distribution of representative st-Xprop domains (domains 3, 4, 8, and 10). g) Domain-specific genes for domains 3, 4, 8, and 10 (from top to bottom). h) GO and KEGG enrichment analysis of differentially expressed genes comparing tumor-related domains (domains 3, 8, and 10) with adjacent normal tissue (domain 4).

In addition, all methods consistently identified the composite lesion formed by DCIS/LCIS3 and IDC 6, although the inferred boundary between these regions deviated slightly from the manual annotation. Boundary locations were nevertheless largely concordant across methods, suggesting shared recognition of intralesional structure. To further interrogate this region, we examined two st-Xprop domains (domains 8 and 12; Fig. 5d) and performed differential expression analysis using the Wilcoxon test (Fig. 5e). These domains exhibited marked transcriptional differences, indicating molecular substructure within a pathologically continuous lesion. Notably, several differentially expressed genes highlighted in Fig. 5e have previously been reported to be aberrantly expressed in breast cancer or associated with tumor aggressiveness and clinical outcome, including immune-related markers (FCGR3B, PSMB9), metabolic regulators (COX6C, DEGS1), and epithelial or secretory genes (MUC5B) [18, 27, 33, 47, 48]. This molecular differentiation aligns with clinical evidence that DCIS can serve as a precursor to IDC, often displaying gradient-like spatial progression [23]. While all compared methods detected partial heterogeneity within this composite lesion, some methods, such as SEDR and STAGATE, produced fragmented subclusters, whereas others, including DeepGFT, SpaGCN, and stLearn, yielded less coherent spatial boundaries. In contrast, st-Xprop delineated the two subregions more coherently, with clearer spatial continuity.

To further characterize intratumoral heterogeneity and assess the biological coherence of st-Xprop domains, we conducted downstream analyses on representative domains (Fig. 5f), including tumor edge (domain 3), healthy tissue (domain 4), DCIS/LCIS (domain 8), and IDC (domain 10). Domain-specific genes were identified using Wilcoxon tests (Fig. 5g). Genes enriched in each domain displayed spatially restricted expression patterns that closely matched the corresponding inferred regions. The tumor edge domain (domain 3) showed higher expression of IGHG1, IGHG3, IGLC2, and IGKC, consistent with enrichment of tumor-infiltrating B cells and plasma cell populations [49]. The healthy tissue domain (domain 4) exhibited increased expression of MGP and S100A6, predominantly localized to non-malignant regions, aligning with known profiles of normal breast epithelium [39, 46]. The DCIS/LCIS domain (domain 8) was characterized by spatially enriched expression of COX6C, CPB1, and CFB, genes previously associated with DCIS progression and inflammatory responses [23, 42]. In contrast, the IDC domain (domain 10) showed higher expression of CXCL14 and CCND1, reflecting enhanced proliferative and invasive activity [26, 29]. Collectively, the concordance between spatially restricted gene expression and known region-specific biological functions supports the anatomical and pathological relevance of the domains identified by st-Xprop.

To further evaluate the biological distinctiveness of the identified tumor-associated domains, we compared tumor-related regions (domains 3, 8, and 10) with the healthy region (domain 4) using differential expression analysis followed by GO and KEGG enrichment (Fig. 5h). Upregulated genes were significantly enriched in extracellular matrix (ECM)–related processes and pathways, including ECM–receptor interaction, cytokine–receptor interaction, chemokine signaling, and hematopoietic lineage, indicating active tumor–stroma crosstalk and immune recruitment. Downregulated genes were enriched in antigen receptor–mediated signaling and pathways such as PI3K–Akt signaling and focal adhesion, suggesting impaired immune surveillance and altered cell adhesion. Notably, ECM–receptor interaction appeared in both up- and downregulated gene sets, reflecting the complex and context-dependent regulation of ECM dynamics within the tumor microenvironment.

Overall, these analyses demonstrate that st-Xprop identifies spatial domains that are not only aligned with histological annotations but also supported by region-specific transcriptional signatures and functional pathway enrichment.

### st-Xprop Identifies Spatially Structured Tumor Programs in a HER2-positive Breast Cancer Dataset

To assess the robustness and generalizability of st-Xprop, we analyzed a multi-patient HER2-positive breast cancer spatial transcriptomics dataset [1], comprising tissue sections from eight patients (groups A–H), with only one annotated section per group. Spatial clustering accuracy was evaluated on annotated sections using the ARI, where Fig. S4 is for sections A-G, Fig. 6a is for section H1. Among the patients and sections evaluated, st-Xprop achieved the highest ARI values, indicating superior cross-slice and cross-patient spatial domain identification.

**Figure 6.**
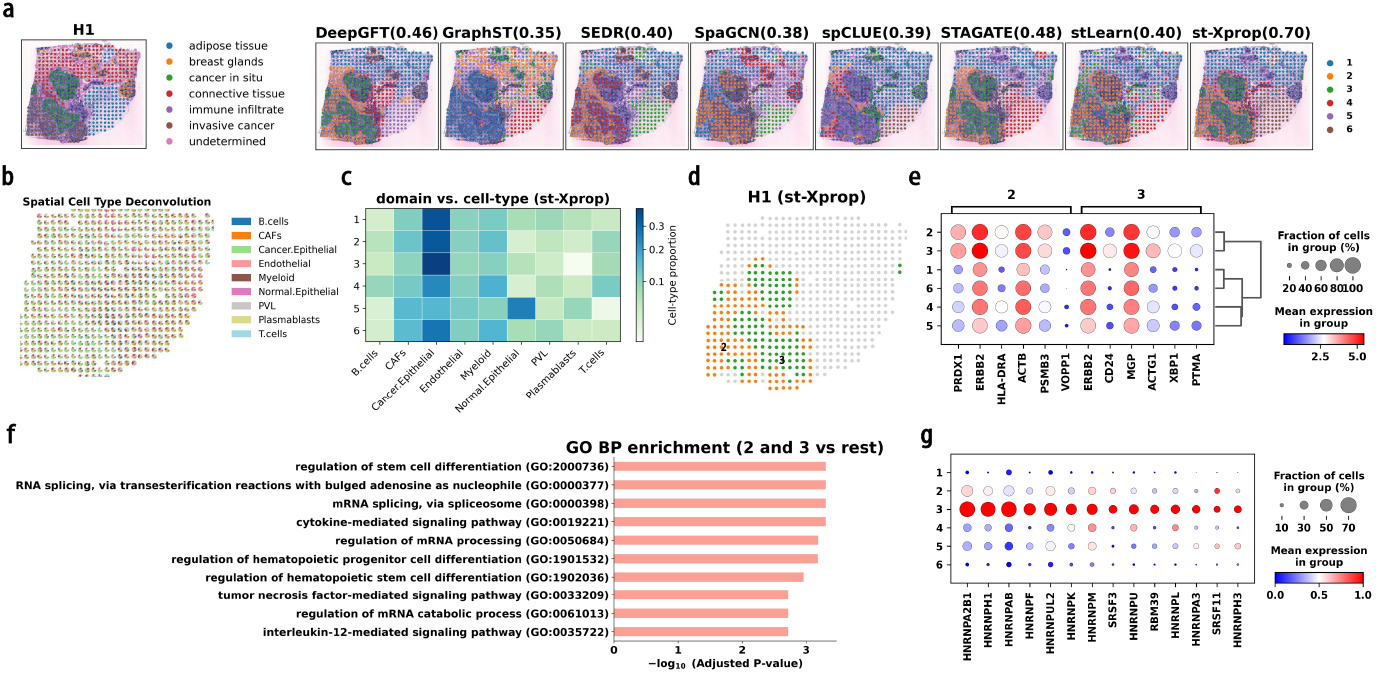
Spatial domain identification in a HER2-positive breast cancer dataset. a) Manual spatial annotation, spatial clustering results obtained by different methods, and corresponding adjusted Rand index (ARI) scores in section H of the HER2-positive breast cancer dataset. b) Spatial distribution of cell-type proportions estimated by RCTD in the H1 section. c) Correspondence between spatial domains identified by st-Xprop and RCTD-inferred cell-type compositions, illustrating domain–cell type concordance. d) Spatial distribution of domains 2 and 3 identified by st-Xprop. e) Top six differentially expressed genes for domains 2 and 3 compared with all other spatial domains. f) GO enrichment analysis of differentially expressed genes in domains 2 and 3 relative to other regions. g) Differentially expressed genes in domains 2 and 3 associated with RNA splicing– and mRNA processing–related GO terms.

We focused subsequent analyses on the H1 section, which exhibits the most diverse tissue architecture. As shown in Fig. 6a, although all methods successfully distinguished cancerous from non-cancerous regions, their performance differed in finer structural resolution. Methods such as GraphST, SpaGCN, spCLUE, and stLearn tended to group in situ and invasive cancer regions into a single cluster. In contrast, DeepGFT, SEDR, STAGATE and st-Xprop were able to separate in situ carcinoma from invasive carcinoma. Notably, st-Xprop generated boundaries that were most consistent with pathological annotations and provided improved delineation of immune-infiltrated and adipose regions. These results suggest that st-Xprop achieves enhanced resolution of fine-grained tissue structures that are often challenging to distinguish in spatial clustering analyses.

To evaluate the biological relevance of the inferred spatial domains, we performed downstream analyses. Using RCTD [5], we estimated spot-level cell-type compositions and visualized the spatial deconvolution results in Fig. 6b. The cancer epithelial cells were broadly distributed across the tissue, with particularly high enrichment in the lower-left region, whereas other cell types predominated in non-cancerous areas, indicating clear spatial heterogeneity in cellular composition.

To further examine the concordance between the identified st-Xprop domains and cellular composition, we constructed a domain–cell-type proportion matrix summarizing the average cell-type proportions within each st-Xprop domain based on the RCTD deconvolution results (Fig. 6c). Distinct cellular compositions were observed across the six inferred domains. Domains 1, 2, and 3 showed strong enrichment of cancer epithelial cells, with domain 3 exhibiting the highest cancer proportion, consistent with their localization to tumor-associated regions. In contrast, domain 5 displayed the lowest cancer cell proportion and was predominantly enriched with normal epithelial cells, corresponding to breast glandular structures and reflecting their largely non-malignant composition. Domain 4 was characterized by increased proportions of immune cells (B cells and T cells) and myeloid cells, consistent with an immune-infiltrated tumor microenvironment. Domain 6 was associated with adipose tissue and exhibited intermediate levels of cancer epithelial cells. This region was also enriched for myeloid and endothelial cells, suggesting spatial mixing between tumor and adipose compartments at the tumor boundary. These results indicate that the st-Xprop-inferred domains capture spatially coherent regions with distinct cellular compositions, highlighting the ability of st-Xprop to recover biologically meaningful tissue architecture.

We next focused on cancer-associated domains 2 and 3, which were identified as tumor-enriched regions based on the spatial clustering results (Fig. 6d), and performed differential expression analysis between tumor-related regions and non-tumor domains (Fig. 6e). Several genes, including PRDX1, ERBB2, ACTB, PSMB3, and MGP, were consistently upregulated in tumor regions, reflecting shared tumor-associated transcriptional programs. Among these, ERBB2 encodes the HER2 receptor, a well-established oncogenic driver in breast cancer whose amplification and overexpression define HER2-positive disease and guide targeted therapy. Its elevated expression across both carcinoma states is consistent with its central role in tumor growth and maintenance [16]. PRDX1 has been implicated in tumor proliferation and oxidative stress regulation and serves as an independent prognostic marker in estrogen receptor–positive breast cancer [34]. ACTB overexpression is associated with cytoskeletal remodeling and enhanced invasive potential [19], while PSMB3, a proteasome subunit often co-expressed with ERBB2, contributes to increased proteasomal activity and tumor progression [11]. MGP encodes a vitamin K–dependent protein and has been associated with poor prognosis and increased invasiveness in breast cancer [56]. These findings indicate that st-Xprop–identified tumor regions capture biologically meaningful transcriptional signatures associated with malignancy.

To further characterize molecular programs associated with tumor-associated regions, we performed GO enrichment analysis on genes upregulated in domains 2 and 3 relative to the remaining tissue regions (Fig. 6f). Enriched GO terms included stemness-related processes and cytokine-mediated signaling pathways, both of which have been widely implicated in invasive breast cancer and its tumor microenvironment [22]. Notably, several enriched terms were related to post-transcriptional regulation. To determine whether this signal reflected functional regulatory programs rather than statistical noise, we further examined the expression patterns of alternative splicing regulators across domains. Among the genes significantly upregulated in domains 2 and 3, we identified several canonical splicing factors, including HNRNPA2B1, HNRNPM, and SRSF3 [17, 25, 55] (Fig. 6g). These factors are known to regulate RNA splicing and transcriptome plasticity in cancer, supporting the biological relevance of the post-transcriptional regulatory programs enriched in tumor-associated domains.

Collectively, these results demonstrate that st-Xprop robustly generalizes across patients and tissue sections, accurately identifies spatially coherent tumor regions, and enables mechanistic insights into cellular composition, transcriptional programs, and regulatory heterogeneity in HER2-positive breast cancer.

### st-Xprop Reveals Coherent Spatial Domains in the PDAC-A Dataset

We further evaluated st-Xprop on a microarray-based spatial transcriptomics dataset from human pancreatic ductal adenocarcinoma (PDAC), the PDAC-A dataset [32]. This dataset presents a particular challenge because ductal, pancreatic epithelial, and stromal compartments are spatially interwoven in PDAC tissues, resulting in gradual transitions rather than sharply defined boundaries. In addition, the limited resolution of microarray-based platforms further complicates the separation of adjacent tissue compartments.

We performed spatial domain identification using different methods (Fig. 7a). Manual annotations, clustering results, and corresponding ARI scores are shown in Fig. 7a. While most methods reliably distinguished tumor and ductal epithelial regions, pancreatic epithelial and stromal compartments appeared more spatially dispersed and were difficult to resolve consistently. In contrast, st-Xprop produced more spatially coherent domains and achieved the highest ARI, indicating improved agreement with expert annotations and enhanced robustness in resolving mixed or transitional regions.

**Figure 7.**
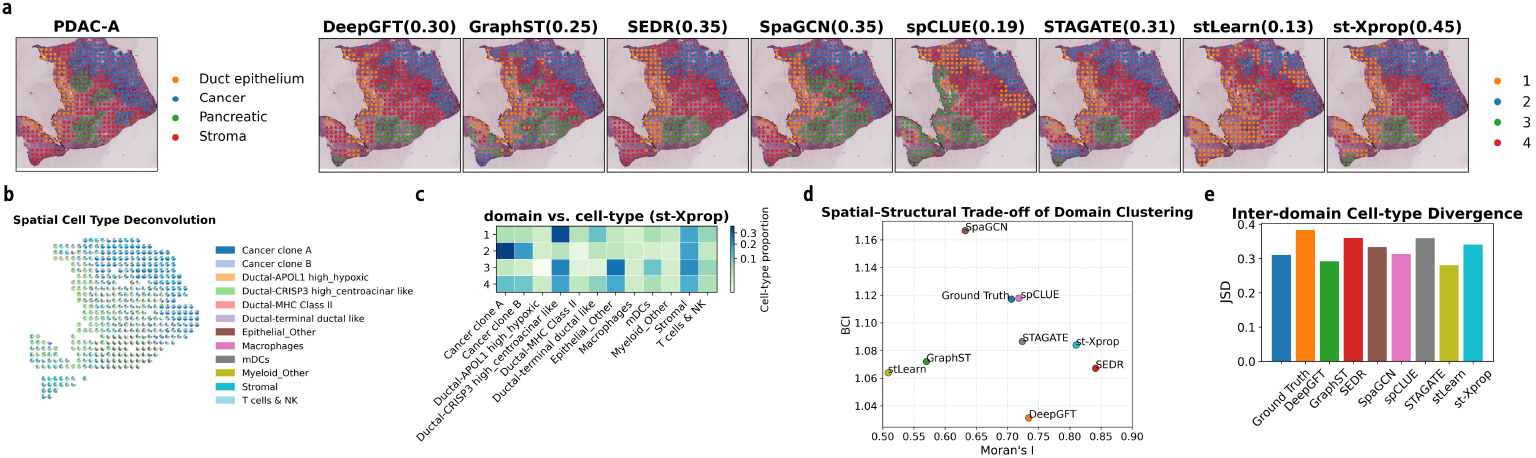
Spatial domain identification results on the PDAC-A dataset. a) Manual spatial domain annotations of the PDAC-A tissue section, clustering results obtained by different methods, and the corresponding ARI scores. b) Spatial distribution of cell-type proportions estimated by RCTD. c) Correspondence between spatial domains identified by st-Xprop and RCTD-inferred cell-type compositions, illustrating domain–cell type concordance. d) Joint evaluation of spatial domains using Moran’s I and the boundary consistency index (BCI), assessing spatial continuity and embedding-level separation, respectively. e) Jensen–Shannon divergence (JSD) between spatial domains based on inferred cell-type compositions, quantifying compositional distinctiveness.

To further interpret the identified domains, we inferred spot-level cell-type composition using RCTD [5]. The inferred cell-type proportions for each spot are shown in Fig. 7b. To systematically evaluate the relationship between spatial domains and cellular composition, we constructed a domain–cell-type proportion matrix summarizing the average cell-type proportions within each st-Xprop domain (Fig. 7c). As illustrated in these analyses, each st-Xprop domain exhibited a distinct and biologically interpretable cellular profile. Domain 1, corresponding to the ductal epithelium region, was enriched for Ductal-CRISP3 high centroacinar-like cells, consistent with a duct-associated niche. Domain 2, corresponding to the cancer region, showed the highest proportion of cancer clone A cells, together with cancer clone B cells, reflecting a tumor-enriched microenvironment. Domain 3, largely aligned with stromal regions, was enriched for stromal components (including endothelial and fibroblast cells) together with epithelial_other cell types (including acinar, endocrine, and tuft cells).

This pattern reflects the well-known interdigitation of epithelial and stromal compartments in PDAC, and suggests that domain 3 captures a biologically heterogeneous microenvironment. Domain 4, corresponding to a duct–tumor interface region, displayed mixed enrichment of Ductal-CRISP3 high centroacinar-like cells and cancer clone B cells, indicating a transitional zone between ductal and malignant compartments. In addition, domains 3 and 4 exhibited partial overlap in inferred cell-type composition. This overlap likely reflects the intrinsic biological intermixing of epithelial and stromal components in PDAC, as well as the limited spatial resolution of microarray-based platforms, where individual spots may capture multiple cell types. Notably, although domains 1 and 2 identified by st-Xprop closely matched the manually annotated structures, domains 3 and 4 partially deviated from the ground truth. However, spots within domains 3 and 4 exhibited relatively consistent cell-type composition profiles in the RCTD results (Fig. 7b). This observation suggests that, while manual annotations primarily reflect morphological boundaries, the st-Xprop-defined domains may better capture transcriptionally defined cellular organization and reveal biologically meaningful heterogeneity. These results indicate that st-Xprop not only achieves high clustering accuracy but also identifies spatial domains that are biologically coherent, even in complex tissues characterized by mixed cell populations and gradual structural transitions.

To further systematically compare the quality of spatial domains identified by different methods, we evaluated clustering results using three complementary metrics: Moran’s I (MI) to quantify spatial continuity, a boundary consistency index (BCI) to assess domain separation in the learned embedding space, and Jensen–Shannon divergence (JSD) to measure differences in cell-type composition between domains (Supplementary file). Higher values of these metrics indicate stronger spatial coherence, clearer embedding-level separation, and greater compositional specificity, respectively. As shown in Fig. 7d, all methods achieved BCI values greater than 1, indicating that inter-domain differences exceeded intra-domain variability in the embedding space. SEDR (MI = 0.841) and st-Xprop (MI = 0.810) achieved the highest Moran’s I scores, indicating strong spatial coherence, whereas GraphST and stLearn exhibited substantially lower MI values, reflecting more fragmented or interleaved domains. Notably, manual annotations did not maximize Moran’s I, consistent with the presence of thin layers, curved boundaries, or interdigitated regions in real tissue that have anatomical significance but reduce global spatial autocorrelation.

In terms of compositional distinctiveness (Fig. 7e), DeepGFT, SEDR, STAGATE, and st-Xprop achieved relatively higher JSD values compared with the other methods. Because JSD measures the divergence between cell-type composition distributions across domains, higher values indicate that the inferred spatial domains are associated with more distinct cellular compositions. These results suggest that the domains identified by these methods better capture biologically differentiated cellular environments within the tissue. Taken together with the spatial continuity and boundary metrics described above, st-Xprop demonstrates a favorable balance between spatial coherence and biologically meaningful compositional differentiation. Notably, manual annotations do not dominate any single quantitative metric. This observation highlights the intrinsic complexity of PDAC tissue architecture and suggests that morphology-based annotations alone may not fully capture the transcriptional and cellular heterogeneity revealed by spatial transcriptomics.

## Discussion

Accurate delineation of spatial domains is fundamental to understanding tissue architecture and its associated biological functions. In this work, we introduce st-Xprop, a cross-propagative learning framework that integrates gene expression profiles, spatial coordinates with histological image features for spatial transcriptomics analysis. By coupling cross-modal alternating propagation with spatially informed graph learning, st-Xprop learns a low-dimensional representation that supports the robust identification of spatially coherent and biologically interpretable tissue domains.

Across multiple real-world spatial transcriptomics datasets, st-Xprop consistently demonstrates strong performance in spatial domain identification. In particular, it shows enhanced stability and sensitivity in resolving domains characterized by weak transcriptional signals, small spatial extent, or complex boundaries. Comparative analyses and ablation experiments further underscore the importance of jointly modeling cross-modal information and spatial structure, highlighting that neither modality alone is sufficient to achieve reliable fine-grained domain segmentation.

Beyond performance improvements, st-Xprop offers a general and extensible framework for spatial domain modeling. While this study primarily focuses on spatial domain identification within individual tissue sections, several promising directions remain for future work. The graph-based formulation naturally lends itself to extension across multiple tissue sections or temporally ordered samples, enabling the study of spatial domain dynamics. In addition, the cross-modal propagation mechanism provides a flexible interface for incorporating additional spatial modalities, such as protein abundance or chromatin accessibility, which may further enhance domain resolution. Finally, integrating weak anatomical supervision or coupling learned domain representations with downstream functional analyses could strengthen biological interpretability and facilitate mechanistic insights into spatial tissue organization.

## Experimental Section

### Datasets and Data Preprocessing

We conducted evaluations on six spatial transcriptomics datasets spanning different scales and species. The basic information of these datasets, including the number of spatial domains and spatial spots, is summarized in Table S1 (Supporting Information). The input to st-Xprop consists of a gene expression matrix **X**_0_ ∈ℝ^*n*×*m*^, spatial coordinates **P** ∈ℝ^*n*×2^, and the corresponding histological image for each tissue section.

Gene expression preprocessing was performed using the `Scanpy` package [51]. Raw count matrices were library-size normalized to a target sum of 10^4^ per spot and log-transformed. Highly variable genes were then selected using the `filter_genes_dispersion` function, and the top 3,000 genes were retained for downstream analysis. Principal component analysis (PCA) was subsequently applied to obtain a 50-dimensional low-dimensional expression representation **X**_*E*_ ∈ℝ^*n*×50^.

Morphological features were extracted from histological images using either stMVC [59] or a pre-trained Vision Transformer (ViT) [7]. Specifically, stMVC employs multi-view contrastive learning to generate spot-level morphological embeddings. For the ViT-based approach, fixed-size image patches (e.g., 224 × 224 pixels) centered at each spot were cropped and processed, with feature representations obtained from the [CLS] token or by averaging patch token embeddings. The resulting image features were assembled into an image feature matrix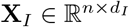.

### Construction of the Spatial Graph and Histological Graph

In st-Xprop, graph structures are constructed from two complementary perspectives: spatial proximity and histological similarity. Specifically, we define two graphs, **G**_*P*_ = (**A**_*P*_, **X**_*E*_) and **G**_*I*_ = (**A**_*I*_, **X**_*E*_), where nodes correspond to spatial spots and node features are given by the low-dimensional gene expression representation **X**_*E*_. The spatial graph **G**_*P*_ encodes local neighborhood relationships based on physical coordinates. Its adjacency matrix **A**_*P*_ is constructed using a radius-based neighbor strategy, such that edges connect spatially proximal spots within a predefined distance threshold, thereby capturing the intrinsic spatial continuity of tissue architecture. In parallel, the histological graph **G**_*I*_ captures morphological similarity between spots. Its adjacency matrix **A**_*I*_ is derived by applying a *k*-nearest neighbor (KNN) algorithm to the image feature matrix **X**_*I*_, linking spots with similar histological appearance regardless of their spatial distance.

Although the two graphs share the same set of nodes and node features, their edge structures differ, reflecting complementary spatial and morphological relationships. Together, these dual graph constructions provide the structural basis for subsequent cross-modal propagation module in st-Xprop.

### Cross-modal Graph Network

The core component of st-Xprop is a cross-modal graph autoencoder that enables effective information exchange between spatial proximity and histological similarity, with the goal of learning robust and discriminative spot-level representations.

We first construct two parallel graph convolutional branches on the spatial and histological graphs to obtain modality-specific embeddings. Given the spatial and image-derived adjacency matrices **A**_*P*_ and **A**_*I*_, we computed their normalized forms using non-symmetric normalization without self-loops:

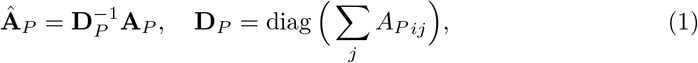

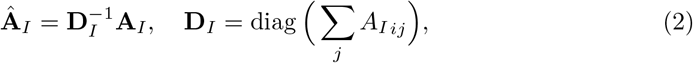

where **D**_*P*_ and **D**_*I*_ are the degree matrices of **A**_*P*_ and **A**_*I*_, respectively. The initial node embeddings for the graph neural network were then computed as

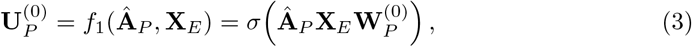

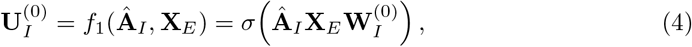

Where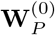and 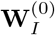 are learnable weight matrices and *σ*(·) denotes the ReLU activation function. These embeddings capture spatially and morphologically informed contextual features, respectively.

### Cross-propagative Learning Module

To enable cross-modal interaction, a cross-propagative mechanism is employed, wherein node embeddings from one modality are propagated along the adjacency structure of the other modality. Specifically, the embeddings are updated as

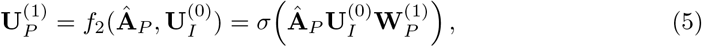

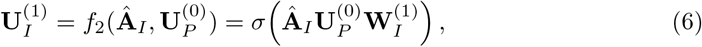

where 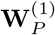 and 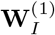 are layer-specific trainable parameters. This alternating propagation allows spatial relationships to modulate histology-informed features and vice versa, thereby coupling complementary structural cues across modalities. The resulting graph-level embedding is obtained by averaging the two branches:

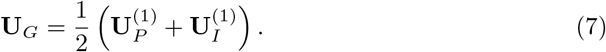

In parallel, gene expression features are independently encoded using a three-layer multilayer perceptron (MLP),

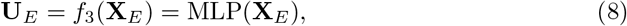

which captures non-linear transcriptional patterns that are complementary to graph-based representations.

The graph-based and expression-based embeddings are then linearly fused to form a unified cross-modal representation:

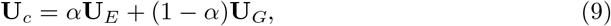

where *α* is a learnable scalar parameter initialized to 0.5, allowing the model to adaptively balance transcriptomic and graph-derived information.

To further capture higher-order structural dependencies, we refine **U**_*c*_ using a lightweight propagation module inspired by DFCN [45]. Specifically, we compute

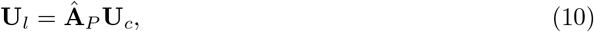

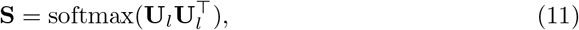

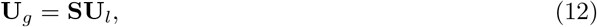

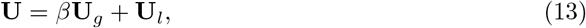

where **U**_*l*_ captures local spatial consistency, **U**_*g*_ encodes global relational semantics, and *β* controls their relative contribution.

### Reconstruction Module

A three-layer fully connected decoder reconstructs the gene expression representation from the latent embedding **U**,

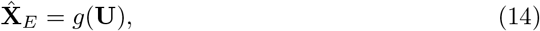

thereby completing the graph autoencoder framework and enabling end-to-end optimization. The decoder is trained by minimizing the reconstruction error:

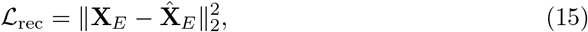

This ensures that the latent representation **U** retains key gene expression information while fusing spatial neighborhood and histological image features, providing a reliable representation for spatial domain identification.

### Clustering Module

After obtaining the fused latent representation **U** from the cross-modal graph network, spatial domains are identified via a self-supervised clustering framework augmented with spatial regularization. This module integrates three complementary components: (i) self-optimizing clustering based on Deep Embedded Clustering (DEC) [52], (ii) spatial smoothing to enforce local coherence, and (iii) a minimum cut constraint to prevent fragmented clusters.

### Self-optimizing Clustering

DEC iteratively refines the embedding space through a procedure of soft assignment, target distribution recalibration, and joint update of embeddings and cluster centers, promoting compact and well-separated clusters. To initialize the clustering, K-means is applied to **U** to obtain *C* initial cluster centroids 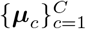. Soft assignment probabilities between spots and centroids are computed using the Student’s *t*-distribution:

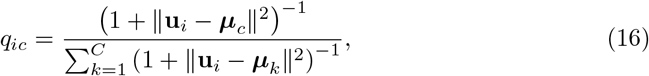

where **u**_*i*_ denotes the embedding of spot *i* and ***μ***_*c*_ is the centroid of cluster *c*. To emphasize high-confidence assignments, an enhanced target distribution is defined as

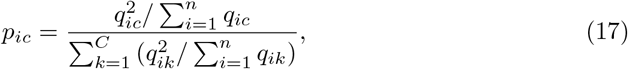

which sharpens the embeddings towards their most likely clusters. The clustering objective is then formulated as the Kullback–Leibler (KL) divergence between the target and soft assignment distributions:

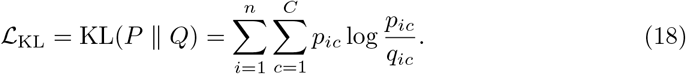

### Spatial Smoothing

To encourage spatial continuity, adjacent spots are regularized to have similar cluster assignments. Let **q**_*i*_ denote the soft assignment vector for spot *i* and **A** the adjacency matrix. The spatial smoothing loss is defined as:

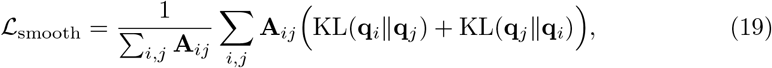

where the adjacency matrix **A** depends on the image feature type:

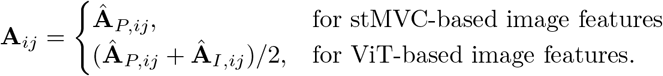

### Minimum Cut Regularization

To prevent cluster fragmentation and enforce global cluster coherence, a minimum cut loss penalizes inconsistent assignments among neighboring spots:

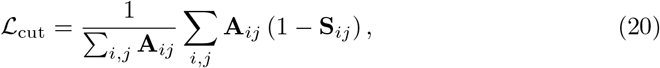

where 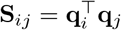 represents pairwise cluster similarity.

This module integrates self-optimizing clustering with spatial smoothing and minimum cut regularization to encourage clusters that exhibit transcriptional similarity and spatial continuity, facilitating biologically interpretable tissue segmentation.

### Loss Function

The st-Xprop model is trained end-to-end by minimizing a composite loss that jointly balances reconstruction fidelity, clustering consistency, and spatial regularization:

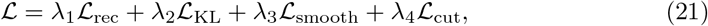

where the first term ensures that the latent representation preserves the key gene expression information, while the second term enforces alignment between soft cluster assignments and the enhanced target distribution, promoting compact and separable clusters. The remaining terms, ℒ_smooth_ and ℒ_cut_, incorporate spatial priors to encourage neighboring spots to have similar assignments and reduce cluster fragmentation. Hyperparameters *λ*_1_–*λ*_4_ control the relative contributions of each component, with default values set to *λ*_1_ = 2, *λ*_2_ = 10, *λ*_3_ = 2, and *λ*_4_ = 1.

### Model Training and Optimization

st-Xprop is trained using a two-stage pretraining–clustering strategy to stabilize representation learning and reduce potential instability from direct end-to-end optimization. During pretraining, the cross-modal graph network is optimized solely with the reconstruction loss ℒ_rec_, yielding an initial embedding **U** that integrates gene expression and graph-based features. In the clustering stage, cluster centroids are initialized with K-means, and soft assignments are iteratively refined using the DEC framework. The total loss ℒ is then minimized jointly over the graph network and clustering parameters, progressively tightening intra-cluster distributions, enhancing inter-cluster separation, and preserving spatial coherence.

### Spatial Domain Characterization and Functional Interpretation

Based on the low-dimensional embeddings learned by st-Xprop, we conducted a series of downstream analyses to characterize spatial domains and investigate their functional and interactional properties.

### Spatial Domain Identification

st-Xprop produces a unified low-dimensional representation for each spatial spot by jointly modeling gene expression, spatial location, and histological features through cross-modal graph propagation. Spatial domains were identified by clustering these embeddings using mclust [38], with the number of clusters informed by reference anatomical annotations.

To functionally characterize each spatial domain, differential expression analysis was performed using the Wilcoxon rank-sum test, comparing spots within a given domain against all remaining spots. For each domain, the top *n*_top_ genes ranked by statistical significance were selected for downstream analysis. These genes were mapped to NCBI Entrez identifiers and subjected to Gene Ontology (GO) and KEGG pathway enrichment analyses using GSEApy [14]. Enrichment was performed using the species-specific genome as background, with multiple testing correction applied via the Benjamini–Hochberg procedure [2]. Pathways with a false discovery rate (FDR) below were considered statistically significant.

### Cell–Cell Interaction Network Construction

To investigate spatially resolved intercellular communication patterns, we constructed cell–cell interaction (CCI) networks based on st-Xprop embeddings following spot-level cell type annotation. Candidate ligand–receptor (LR) pairs were curated from publicly available databases [8, 12] and used as potential communication events.

Given that each spatial spot may contain multiple cells, LR pairs were first evaluated at the spot level. A spot was considered eligible for communication if both a ligand and its corresponding receptor were highly expressed. A *k*-nearest neighbor (kNN) graph was then constructed using st-Xprop embeddings to define spatially informed neighborhood relationships between spots. If a spot annotated as cell type A expressed ligand *L* and a neighboring spot annotated as cell type B expressed the corresponding receptor *R*, this was counted as an *A*→ *B* communication event. Aggregation over all spot pairs yielded a weighted cell–cell interaction network.

Statistical significance of inferred interactions was assessed using permutation testing. Spatial coordinates were held fixed while cell type annotations were randomly permuted, and communication strengths were recomputed to generate null distributions. Empirical *p*-values were obtained by comparing observed interaction strengths against the null, and interactions with *p <* 0.05 were retained.

The resulting CCI network was represented as an undirected weighted graph and partitioned into modules using the Leiden algorithm. To assess signaling specificity within each module, ligand–receptor enrichment analysis was performed. Weighted edge counts were used to compute global and module-specific LR usage frequencies, from which 2 × 2 contingency tables were constructed. One-sided Fisher’s exact tests were applied, followed by Benjamini–Hochberg correction. Ligand–receptor pairs with FDR *<* 0.05 were considered significantly enriched within the corresponding module.

### Baseline Methods

We compared st-Xprop with seven representative state-of-the-art methods for spatial domain identification. Among them, SpaGCN [20] and stLearn [35] explicitly incorporate histology-derived features, while the remaining methods rely on gene expression and spatial information. Brief descriptions of the compared methods are provided below.

DeepGFT [41] projects spatial transcriptomics data into the frequency domain defined by two graph structures: a spatial proximity graph among spots and a gene co-expression graph. Graph Fourier transform (GFT) is applied for low-pass filtering to reduce dropout and noise. Dual-channel embeddings are learned using graph attention autoencoders and subsequently fused through inverse transforms and attention mechanisms to reconstruct denoised gene expression, which is then used for spatial domain identification.

GraphST [54] is a self-supervised contrastive learning framework based on graph neural networks. It integrates gene expression profiles and spatial coordinates by constructing a spatial graph and learns spot representations by maximizing the similarity of embeddings for spatially adjacent spots while minimizing that of non-adjacent spots. The learned embeddings are used for downstream spatial clustering.

SEDR [53] is a masked self-supervised variational graph autoencoder framework. It first learns a low-dimensional latent representation of gene expression using a masked autoencoder. Spatial neighborhood information is then incorporated via a variational graph autoencoder, and the resulting spatial embeddings are concatenated with gene representations to form a joint latent space. The model is trained by reconstructing masked gene expression for spatial clustering.

SpaGCN [20] integrates gene expression, spatial coordinates, and histological image features to construct a weighted undirected graph, where edge weights reflect both spatial proximity and histological similarity. Graph convolutional layers aggregate gene expression from neighboring spots, and the resulting representations are clustered using a deep embedding clustering framework to identify spatial domains.

spCLUE [50] is a multi-view graph contrastive learning framework designed for both single-slice and multi-slice spatial transcriptomics analysis. For each slice, it constructs a spatial neighborhood graph based on Euclidean distance and an expression similarity graph based on Pearson correlation. A GCN encoder extracts embeddings from both views, which are aligned using instance-level and cluster-level contrastive learning. An attention mechanism is then used to fuse the dual-view embeddings into a unified representation for spatial domain identification.

STAGATE [10] constructs a spatial adjacency graph based on spot distances, and employs a graph attention network (GAT) to aggregate gene expression from neighboring spots with adaptive attention weights. A decoder reconstructs the original expression profiles, and the model is jointly optimized using reconstruction and clustering objectives. An adaptive edge pruning strategy is applied to balance spatial proximity and expression similarity in the learned embeddings.

stLearn [35] constructs a composite weighted neighborhood graph by integrating spatial coordinates, gene expression profiles, and tissue morphology features extracted from H&E images using a pretrained ResNet. Weighted neighborhood smoothing is performed based on a composite similarity measure, enabling expression denoising and downstream spatial domain analysis while preserving tissue structure.

## Supporting Information

Supporting Information is available in the Supplementary file.pdf.

## Acknowledgments

This work was funded by National Natural Science Foundation of China projects under Grant No. 12222115 and 92470106.

