## Supplementary Figure 1-4, Supplementary Table 1 for "Cross-Propagative Graph Learning Reveals Spatial Tissue Domains in Multi-Modal Spatial Transcriptomics"

### 1 Evaluation Metrics for Spatial Domain Quality

To quantitatively assess the quality of inferred spatial domains, we employed three complementary metrics that capture distinct aspects of spatial organization and biological interpretability: Moran's I (MI) for spatial autocorrelation, the Boundary Coherence Index (BCI) for boundary sharpness in the latent embedding space, and Jensen–Shannon divergence (JSD) for the distinctness of cell-type compositions between domains.

Consider a spatial transcriptomics dataset containing (N) spots. Each spot $i$ is assigned to a spatial domain $k_i \in 1, \ldots, K$ and is associated with a learned low-dimensional embedding $\mathbf{u}_i \in \mathbb{R}^d$. Spatial neighborhood relationships are defined using a kNN graph constructed from spatial coordinates.

#### 1.1 Moran's I (MI)

Moran's I quantifies the degree of spatial autocorrelation of domain assignments over the spatial neighborhood graph. For each spatial domain $k$, we define a binary indicator variable:

$$y_i^{(k)} = \begin{cases} 1, & k_i = k, \\ 0, & \text{otherwise}, \end{cases} \qquad \bar{y}_k = \frac{1}{N} \sum_{i=1}^{N} y_i^{(k)}.$$

Let $W$ denote the spatial weight matrix derived from the kNN graph:

$$w_{ij} = \begin{cases} 1, & j \in kNN(i), \\ 0, & \text{otherwise}, \end{cases}$$

The Moran's I score for domain $k$ is defined as:

$$MI_k = \frac{N}{\sum_i \sum_j w_{ij}} \cdot \frac{\sum_i \sum_j w_{ij} \big(y_i^{(k)} - \bar{y}_k\big)\big(y_j^{(k)} - \bar{y}_k\big)}{\sum_i \big(y_i^{(k)} - \bar{y}_k\big)^2}.$$

The overall spatial autocorrelation score is computed by averaging over all domains:

$$\text{MI} = \frac{1}{K} \sum_{k=1}^{K} \text{MI}_k.$$

Positive MI values indicate spatial clustering of domain labels, values close to zero correspond to spatial randomness, and negative values indicate spatial dispersion. MI therefore measures the spatial continuity and coherence of inferred domains.

#### 1.2 Boundary Coherence Index (BCI)

The Boundary Coherence Index evaluates how well spatial domain boundaries are separated in the learned embedding space. Let the spatial neighborhood graph be represented as an undirected graph:

$$G = (V, E), \quad |V| = N,$$

where $E$ denotes the set of adjacent spot pairs. Edges are partitioned into within-domain edges and between-domain edges:

$$E_b = \{(i, j) \in E \mid k_i \neq k_j\}, \quad E_w = \{(i, j) \in E \mid k_i = k_j\}.$$

We define the average embedding distances for within-domain and boundary edges as:

$$D_b = \frac{1}{|E_b|} \sum_{(i,j)\in E_b} ||\mathbf{u}_i - \mathbf{u}_j||_2, \quad D_w = \frac{1}{|E_w|} \sum_{(i,j)\in E_w} ||\mathbf{u}_i - \mathbf{u}_j||_2.$$

The Boundary Coherence Index is then defined as:

$$\text{BCI} = \frac{D_b}{D_w}.$$

A BCI value greater than 1 indicates that embedding differences across domain boundaries are larger than those within domains, corresponding to sharper and more coherent boundaries in the latent space.

#### 1.3 Jensen–Shannon Divergence (JSD)

To quantify differences in cell-type composition between spatial domains, we first aggregated spot-level cell-type proportions inferred by Tangram into domain-level compositions. Specifically, let $\mathbf{t}_i = (t_{i1}, \ldots, t_{iC})$ denote the cell-type proportion vector for spot $i$, where $t_{ic}$ represents the Tangram-inferred abundance of cell type $c$ at spot $i$, and $C$ denotes the number of reference cell types.

For each spatial domain $k$, we computed the average cell-type composition by aggregating all spots assigned to that domain:

$$\tilde{\mathbf{p}}_k = \frac{1}{|\mathcal{S}_k|} \sum_{i \in \mathcal{S}_k} \mathbf{t}_i,$$

where $\mathcal{S}_k = \{i \mid k_i = k\}$ denotes the set of spots in domain $k$. The resulting vector was then normalized to sum to one:

$$\mathbf{p}_k = \frac{\tilde{\mathbf{p}}_k}{\sum_{c=1}^{C} \tilde{p}_{kc}},$$

yielding a probability distribution over cell types for each spatial domain.

For any pair of domains $k$ and $l$, the Jensen–Shannon divergence is defined as:

$$\mathrm{JSD}(\mathbf{p}_k, \mathbf{p}_l) = \frac{1}{2}\mathrm{KL}(\mathbf{p}_k || \mathbf{m}) + \frac{1}{2}\mathrm{KL}(\mathbf{p}_l || \mathbf{m}), \quad \mathbf{m} = \tfrac{1}{2}(\mathbf{p}_k + \mathbf{p}_l),$$

where the Kullback–Leibler divergence is given by:

$$\mathrm{KL}(\mathbf{p} | \mathbf{q}) = \sum_c p_c \log \frac{p_c}{q_c}.$$

The overall JSD score is computed as the average pairwise divergence across all domain pairs:

$$\mathrm{JSD} = \frac{2}{K(K-1)} \sum_{k<l} \mathrm{JSD}(\mathbf{p}_k, \mathbf{p}_l),$$

Higher JSD values indicate stronger differences in cell-type composition between spatial domains, reflecting increased biological specificity and interpretability.

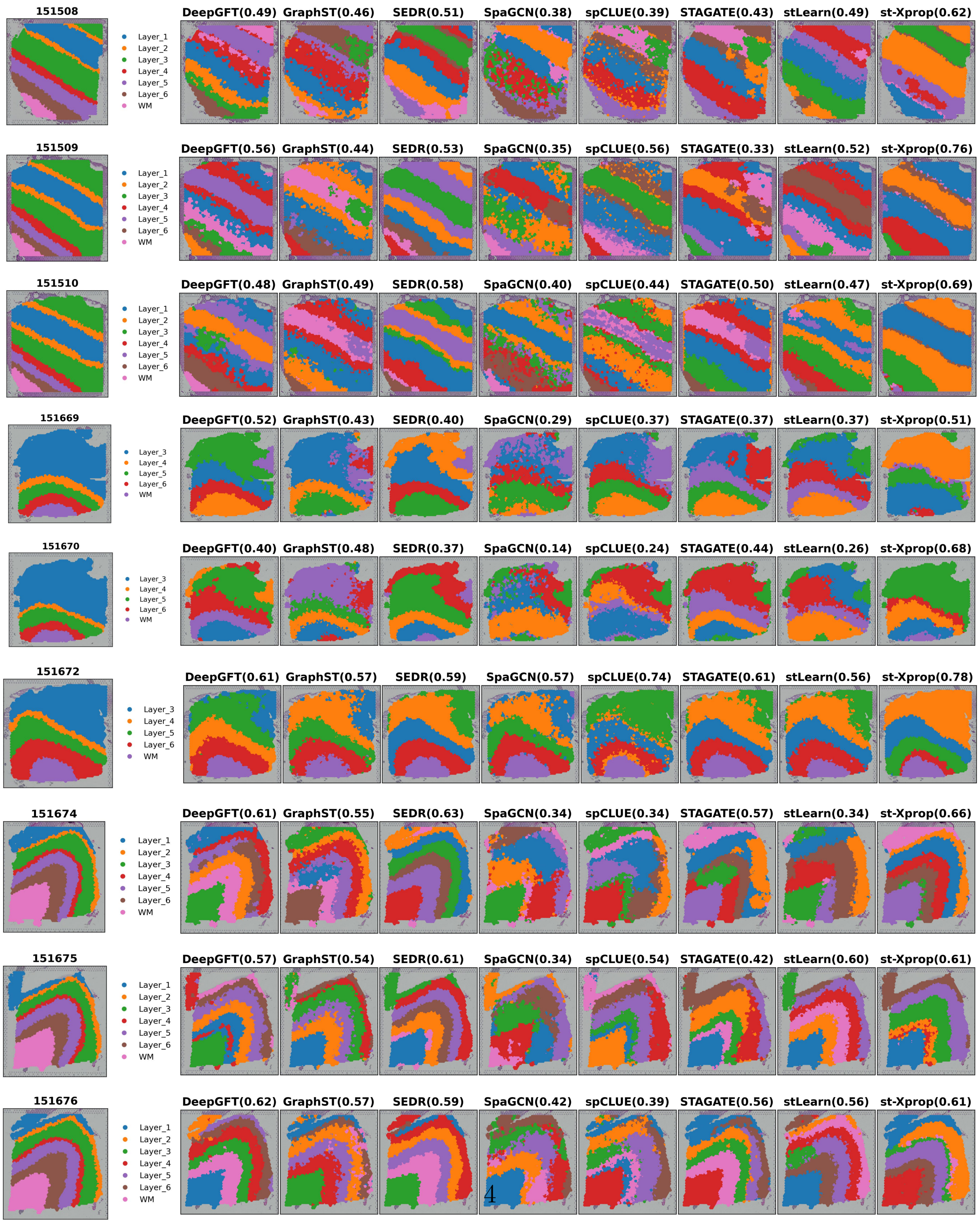




Fig. S1: Comparison of manually annotated spatial domains with the domains identified by DeepGFT, GraphST, SEDR, SpaGCN, spCLUE, STAGATE, stLearn, and st-Xprop across 9 DLPFC slices.

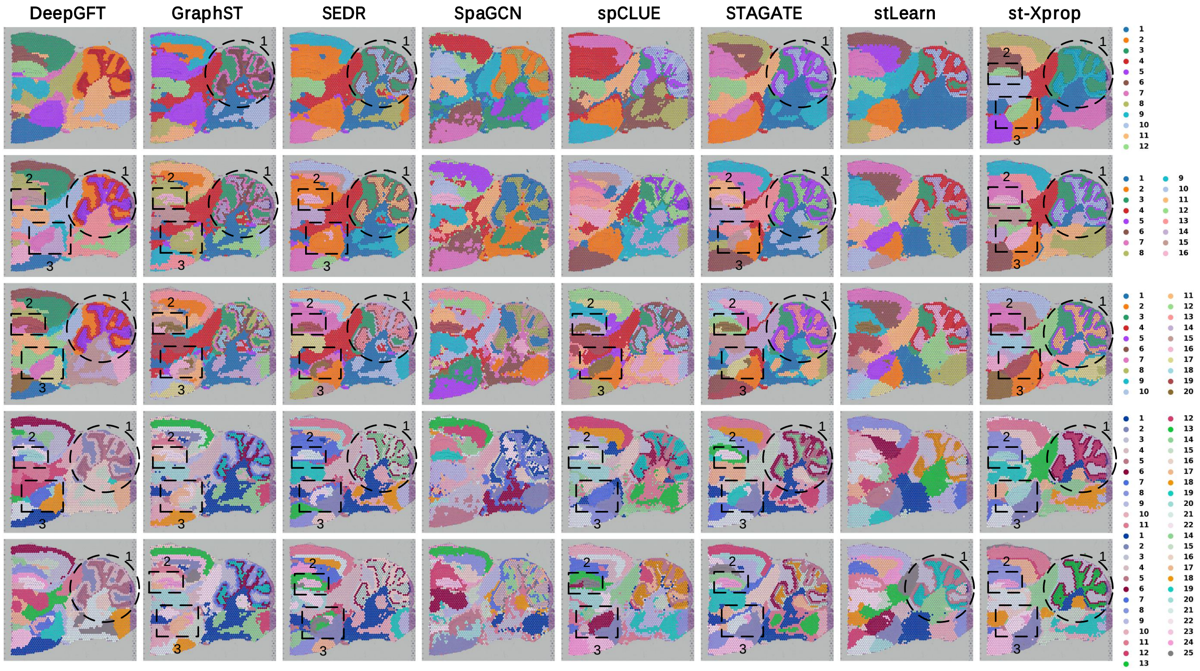


Fig. S2: Spatial domain visualization on the mouse brain posterior (MBP) section identified by different methods. Each row corresponds to a different number of clusters from top to bottom: 12, 16, 20, 22, and 25.

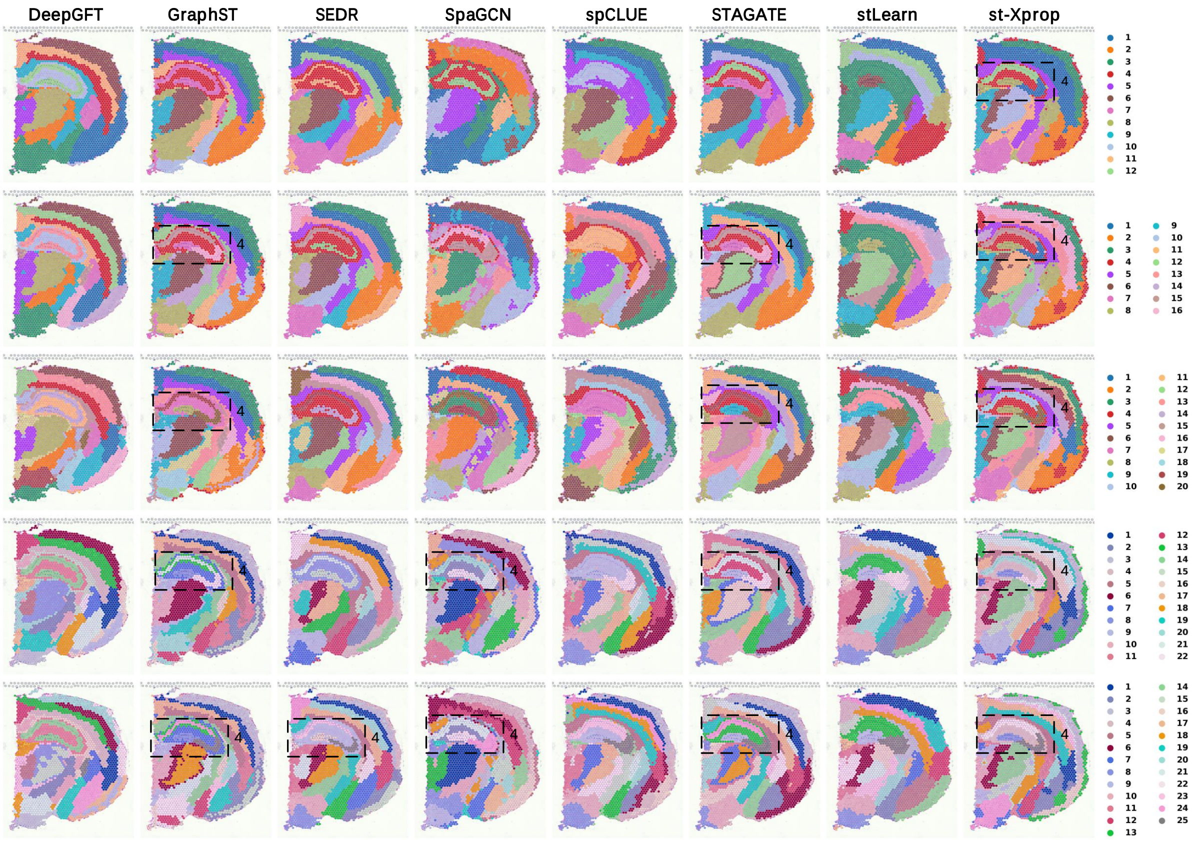


Fig. S3: Spatial domain visualization on the mouse brain coronal (MBC) section identified by different methods. Each row corresponds to a different number of clusters from top to bottom: 12, 16, 20, 22, and 25.

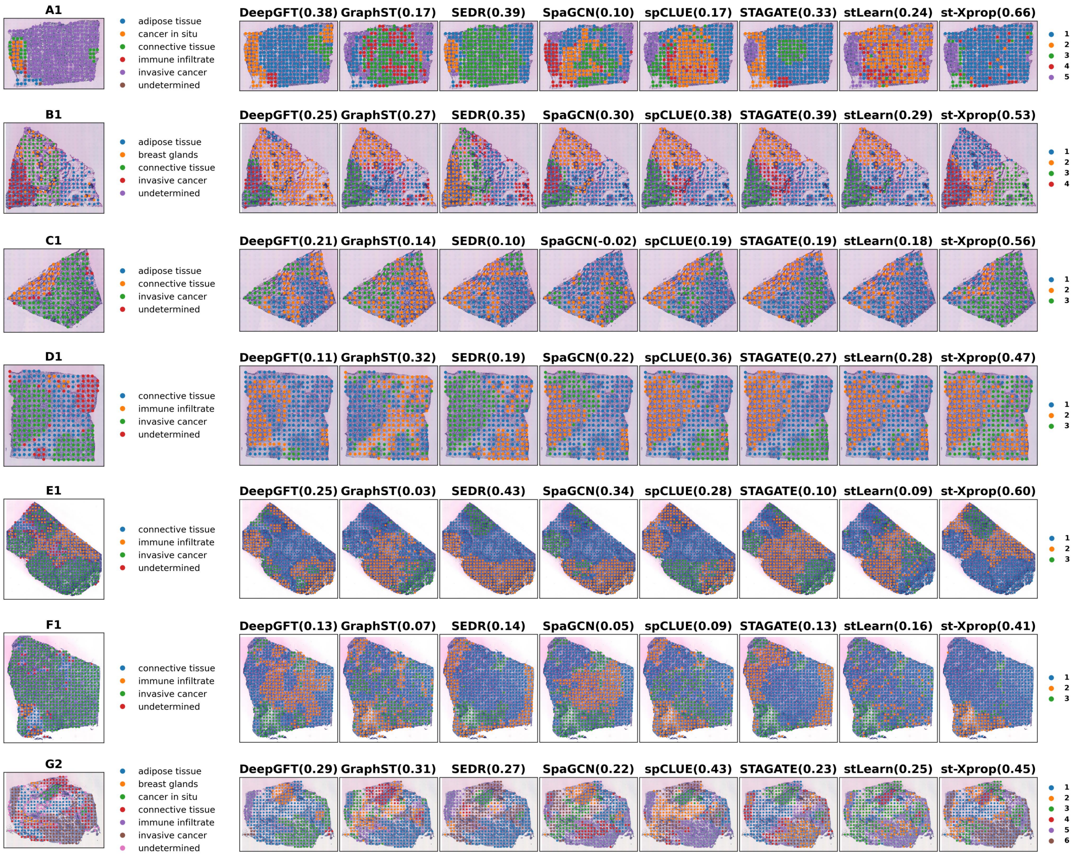


Fig. S4: Manual spatial annotations, spatial clustering results obtained by different methods, and corresponding adjusted Rand index (ARI) scores across sections A-H of the HER2-positive breast cancer dataset.

Table S1: Summary of the ST datasets used in this study.

| Platform | Tissue | Section | # domains | Spots | Related figures |
|---|---|---|---|---|---|
| 10x Visium | Human dorsolateral prefrontal cortex (DLPFC) | 151507 | 7 | 4,226 | Fig. 2 Fig. S1 |
| | | 151508 | 7 | 4,384 | |
| | | 151509 | 7 | 4,789 | |
| | | 151510 | 7 | 4,634 | |
| | | 151669 | 5 | 3,661 | |
| | | 151670 | 5 | 3,498 | |
| | | 151671 | 5 | 4,110 | |
| | | 151672 | 5 | 4,015 | |
| | | 151673 | 7 | 3,639 | |
| | | 151674 | 7 | 3,673 | |
| | | 151675 | 7 | 3,592 | |
| | | 151676 | 7 | 3,460 | |
| 10x Visium | Mouse brain | Sagittal posterior section 1 | 12-25 | 3,355 | Fig. 3 Fig. S2, Fig.S3 |
| | | Coronal section 1 | 12-25 | 2,702 | |
| 10x Visium | Developing chicken heart | D4 | 5 | 747 | Fig. 4 |
| | | D7 | 7 | 1,966 | |
| | | D10 | 7 | 1,916 | |
| | | D14 | 6 | 1,967 | |
| 10x Visium | Human breast cancer | Section 1 | 20 | 3,798 | Fig. 5 |
| 10x Visium | HER2-positive breast cancer | Section A1 | 5 | 346 | Fig. 6 Fig. S4 |
| | | Section B1 | 4 | 295 | |
| | | Section C1 | 3 | 176 | |
| | | Section D1 | 3 | 306 | |
| | | Section E1 | 3 | 587 | |
| | | Section F1 | 3 | 691 | |
| | | Section G2 | 6 | 467 | |
| | | Section H1 | 6 | 613 | |
| ST | Pancreatic ductal adeno-carcinoma | Section A | 4 | 426 | Fig. 7 |
